# The evolutionary fates of a large segmental duplication in mouse

**DOI:** 10.1101/043687

**Authors:** Andrew P Morgan, J Matthew Holt, Rachel C McMullan, Timothy A Bell, Amelia M-F Clayshulte, John P Didion, Liran Yadgary, David Thybert, Duncan T Odom, Paul Flicek, Leonard McMillan, Fernando Pardo-Manuel de Villena

**Affiliations:** Department of Genetics, Carolina Center for Genome Sciences and Lineberger Comprehensive Cancer Center, University of North Carolina, Chapel Hill, NC; Department of Computer Science, University of North Carolina, Chapel Hill, NC; European Molecular Biology Laboratory, European Bioinformatics Institute, Wellcome Genome Campus, Hinxton, Cambridge, CB10 1SD, United Kingdom; University of Cambridge, Cancer Research UK Cambridge Institute, Robinson Way, Cambridge, CB2 0RE, United Kindgom; Wellcome Trust Sanger Institute, Wellcome Genome Campus, Hinxton, Cambridge, CB10 1SA, United Kingdom

**Author notes:** Corresponding author: Fernando Pardo-Manuel de Villena5049 Genetic Medicine Building 120 Mason Farm Road CB#7264 Chapel Hill, NC 27599-7264.

## Abstract

Gene duplication and loss are major sources of genetic polymorphism in populations, and are important forces shaping the evolution of genome content and organization. We have reconstructed the origin and history of a 127 kbp segmental duplication, *R2d*, in the house mouse (*Mus musculus). R2d* contains a single protein-coding gene, *Cwc22. De novo* assembly of both the ancestral (*R2d1*) and the derived (*R2d2*) copies reveals that they have been subject to non-allelic gene conversion events spanning tens of kilobases. *R2d2* is also a hotspot for structural variation: its diploid copy number ranges from zero in the mouse reference genome to more than 80 in wild mice sampled from around the globe. Hemizgyosity for high-copy-number alleles of *R2d2* is associated in *cis* with meiotic drive, suppression of meiotic crossovers, and copy-number instability, with a mutation rate in excess of 1 per 100 transmissions in laboratory populations. We identify an additional 57 loci covering 0.8% of the mouse genome with patterns of sequence variation similar to those at *R2d1* and *R2d2*. Our results provide a striking example of allelic diversity generated by duplication and demonstrate the value of *de novo* assembly in a phylogenetic context for understanding the mutational processes affecting duplicate genes.

## Introduction

Duplication is an important force shaping the évolution of plant and animal genomes: it provides both a substrate for evolution and transient relief from selective pressure (Lynch and Conery 2000). Segmental duplications (SDs), defined as contiguous sequences which map to more than one physical position (Bailey and Eichler 2006), are a common feature of eukaryotic genomes and particularly those of vertebrates.

Like any sequence variant, a duplication first arises in a single individual in a population. The distinction between such copy-number variants (CNVs) and SDs is fluid and somewhat arbitrary: tracts of SDs are highly polymorphic in populations in species from *Drosophila* (Dopman and Hartl 2007) to mouse (She *et al*. 2008) to human (Bailey and Eichler 2006). Studies of parent-offspring transmissions have shown that SDs are prone to recurrent *de novo* mutations including some implicated in human disease (reviewed in Stankiewicz and Lupski 2002). Bursts of segmental duplication have preceded dramatic species radiations in primates. More broadly, blocks of conserved synteny in mammals frequently terminate at SDs (Bailey and Eichler 2006), suggesting that SDs could mediate the chromosomal rearrangements through which karyotypes diverge and reproductive barriers arise.

Notwithstanding their evolutionary importance, SDs are difficult to analyze. Repeated sequences with period longer than the insert size in a sequencing library and high pairwise similarity are likely to be collapsed into a single sequence during genome assembly. Efficient and sensitive alignment of high-throughput sequencing reads to duplicated sequence remains challenging (Treangen and Salzberg 2011). Genotyping of sites within SDs is difficult because variants between copies (paralogous variants) are easily confounded with variants within copies between individuals at a given copy (allelic variants). Latent paralogous variation may bias interpretations of sequence diversity and haplotype structure (Hurles 2002).

Paralogy also complicates phylogenetic inference. Ancestral duplication followed by differential losses along separate lineages may yield a local phylogeny that is discordant with the genome-wide phylogeny (Goodman *et al*. 1979). Within each duplicate copy, local phylogenies for adjacent intervals may also be discordant due to non-allelic gene conversion between copies (Dover 1982; Nagylaki and Petes 1982). As a result, over some fraction of the genome, sequences from individuals of the same species may be more closely related to sequences from individuals of an outgroup species than they are to each other.

In this manuscript we present a detailed analysis of a segmental duplication, *R2d*, in the house mouse (*Mus musculus). R2d* is a 127 kbp unit which contains the protein-coding gene *Cwc22* and flanking intergenic sequence. Although the C57BL/6J reference strain and other classical laboratory strains have a single haploid copy of the *R2d* sequence (in the *R2d1* locus), the wild-derived CAST/EiJ, ZALENDE/EiJ, and WSB/EiJ strains have an additional 1, 16 and 33 haploid copies respectively in the *R2d2* locus. *R2d2* is the responder locus in a recently-described meiotic drive system on mouse chromosome 2 but is absent from the mouse reference genome (Waterston *et al*. 2002; Didion *et al*. 2015, 2016). We draw on a collection of species from the genus *Mus* sampled from around the globe to reconstruct the sequence of events giving rise to the locus’ present structure (**Figure 1**). Using novel computational tools built around indexes of raw high-throughput sequencing reads, we perform local *de novo* assembly of phased haplotypes and explore patterns of sequence diversity within and between copies of *R2d*.

**Figure 1.**
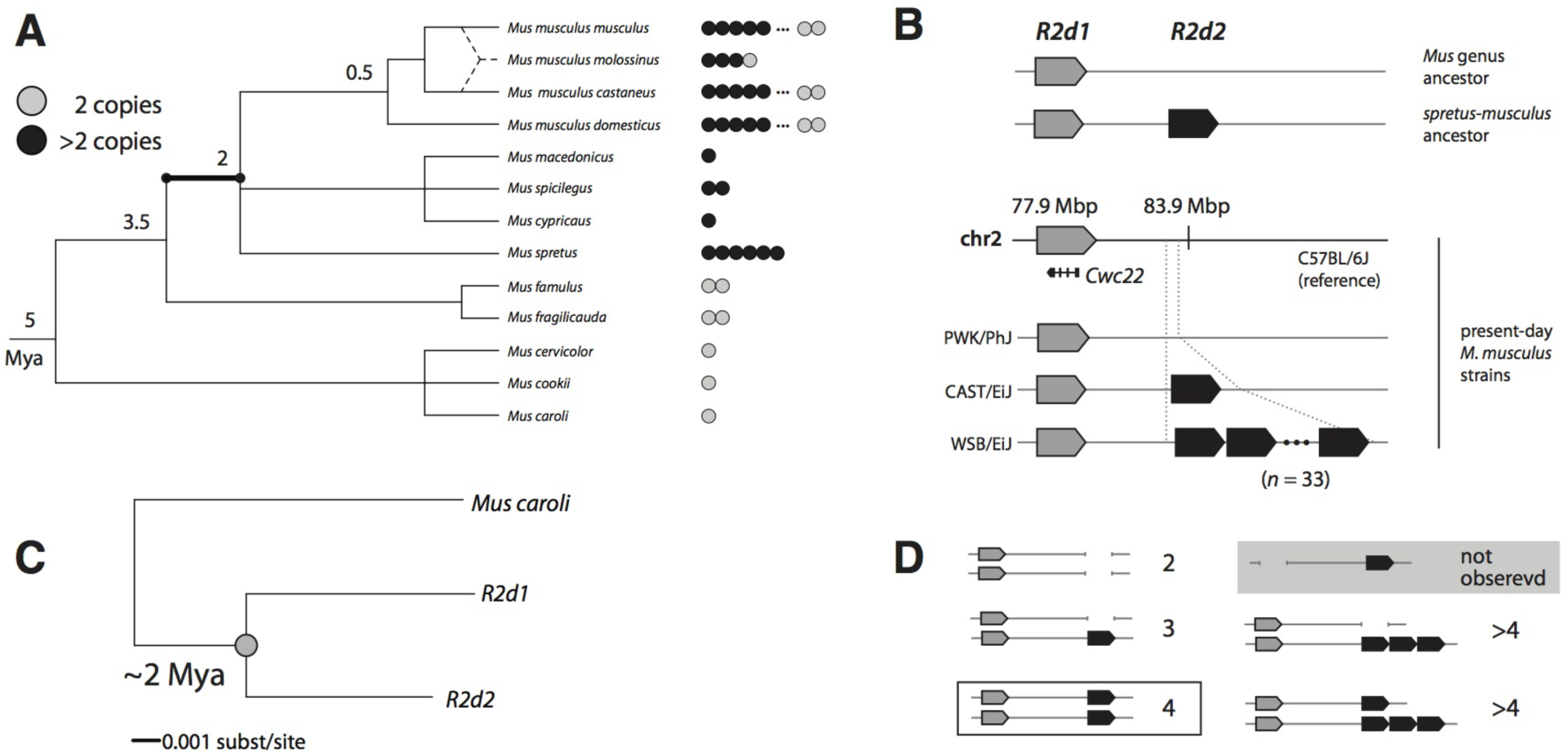
Origin and age of the *R2d2* duplication. **(A)** *Rid* copy number across the phylogeny of the genus *Mus*. Each dot represents one individual; grey dots indicate diploid copy number 2 and black dots copy number >2. The duplication event giving rise to *R2d1* and *R2d2* most likely occurred on the highlighted branch. Approximate divergence times (REF: Suzuki 2004) are given in millions of years ago (Mya) at internal nodes. **(B)** Schematic structure of the *R2d1-R2d2* locus. The mouse reference genome (strain C57BL/6J, *M. m. domesticus*) contains a single copy of *R2d* at *R2d2*. Wild-derived inbred strains vary in haploid copy number from 1 (PWK/PhJ, *M. m. musculus*) to 2 (CAST/EiJ, *M. m. castaneus*) to 33 (WSB/EiJ, *M. m. domesticus*). *R2d2* is located at approximately 77.9 Mbp and *R2d2* at 83.8 Mbp. **(C)** Concatenated tree constructed from *R2d2* (reference genome) and *de novo* assembled *R2d2* and *M. caroli* sequences assuming a strict molecular clock. The duplication node is indicated with a grey dot. **(D)** Relationship between observed *Rid* copy-number states and inferred structure of the *Rldi-Rldl* locus. The configuration of the *M. spretus* - *M. musculus* common ancestor (4 diploid copies) is boxed in black. We have yet to identify samples with diploid copy number 2 but no *R2d1* (grey shaded box).

Both phylogenetic analyses and estimation of mutation rate in laboratory mouse populations reveal that *R2d2* and its surrounding region on chromosome 2 are unstable in copy number. Cycles of duplication, deletion and non-allelic gene conversion have led to complex phylogenetic patterns discordant with species-level relationships within *Mus* which cannot be explained by known patterns of introgression between *Mus* species (Bonhomme *et al*. 2007; Yang *et al*. 2011).

Finally, we identify 57 other loci, covering 0.8% of the mouse genome, which share the key features of *R2d2:* elevated local sequence divergence; low recombination rate; and enrichment for segmental duplications. Previous studies of sequence variation in the mouse (Keane *et al*. 2011) have attributed this pattern to sorting of alleles segregating in the common ancestor of *M. musculus* and its sister species. We suggest instead that these loci have been subject to independent cycles of duplication and loss along *Mus* lineages. Marked enrichment for odorant, pheromone, and antigen-recognition receptors supports a role for balancing selection on the generation and maintenance of the extreme level of polymorphism observed at these loci.

## Results

### *R2d* was duplicated in the common ancestor of *M. musculus* and *M. spretus*

In order to determine when the *R2d* CNV arose, we used quantitative PCR and/or depth of coverage in whole-genome sequencing to assay *R2d* copy number in a collection of samples spanning the phylogeny of the genus *Mus*. Samples were classified as having diploid copy number 2 (two chromosomes each with a single copy of *R2d*) or >2 (at least one chromosome with an *R2d* duplication).

We find evidence for >2 diploid copies in representatives of all mouse taxa tested from the Palearctic clade (Suzuki *et al*. 2004) (**Figure 1** and **Supplementary Table 1**): 236 of 525 *Mus musculus*, 1 of 1 *M. macedonicus*, 1 of 1 *M. spicilegus*, 1 of 1 *M. cypriacus* and 8 of 8 *M. spretus* samples. However, we find no evidence of duplication in species from the southeast Asian clade, which is an outgroup to Palearctic mice: 0 of 2 *M. famulus*, 0 of 2 *M. fragilicauda*, 0 of 1 *M. cervicolor*, 0 of 1 *M. cookii* and 0 of 1 *M. caroli* samples. Outside the subgenus *Mus*, we found evidence for >2 diploid copies in none of the 9 samples tested from subgenus *Pyromys*. We conclude that the *R2d* duplication most likely occurred between the divergence of southeast Asian from Palearctic mice (~3.5 million years ago [Mya]) and the divergence of *M. musculus* from *M. spretus* (~2 Mya) (Suzuki *et al*. 2004; Chevret *et al*. 2005), along the highlighted branch of the phylogeny in **Figure 1A**. If the *R2d* duplication is ancestral to the divergence of *M. musculus*, then extant lineages of house mice which have 2 diploid copies of *R2d* – including the reference strain C57BL/6J (of predominantly *M. musculus domesticus* origin (Yang *et al*. 2007)) – represent subsequent losses of an *R2d* copy.

Duplication of the ancestral *R2d* sequence resulted in two paralogs residing in loci which we denote *R2d1* and *R2d2* (**Figure 1B**). Only one of these is present in the mouse reference genome, at chr2: 77.87 Mbp; the other copy maps approximately 6 Mbp distal (Didion *et al*. 2015), as we describe in more detail below. The more proximal copy, *R2d1*, lies in a region of conserved synteny with rat, rabbit, chimpanzee and human (Muffato *et al*. 2010) (**Supplementary Figure 1**); we conclude that it is the ancestral copy.

The sequence of the *R2d2* paralog was assembled *de novo* from whole-genome sequence reads (Keane *et al*. 2011) from the strain WSB/EiJ (of pure *M. m. domesticus* origin (Yang *et al*. 2011)), which has diploid *R2d* copy number ~68 (Didion *et al*. 2015). We exploited the difference in depth of coverage for *R2d1* (1 haploid copy) and *R2d2* (33 haploid copies) to assign variants to *R2d1* or *R2d2*. Pairwise alignment of the *R2d2* contig against *R2d1* is shown in **Supplementary Figure 2**. The paralogs differ by at least 8 transposable-element (TE) insertions: 7 LINE elements specific to *R2d1* and 1 endogenous retroviral element (ERV) specific to *R2d2* (**Supplementary Table 2**). (Due to the inherent limitations of assembling repetitive elements from short reads, it is likely that we have underestimated the number of young TEs in *R2d2*.) The R2d1-specific LINEs are all < *2%* diverged from the consensus for their respective families in the RepeatMasker database (http://www.repeatmasker.org/cgi-bin/WEBRepeatMasker), consistent with insertion within the last 2 My. The oldest R1d2-specific ERV we could detect is 0.7% diverged from its family consensus. TE insertions occurring since the ancestral *R2d* duplication are almost certainly independent, so these data are consistent with duplication <2 Mya. The *R2d* unit, minus paralog-specific TE insertions, is 127 kbp in size. *R2d* units in the *R2d2* locus are capped on both ends by (CTCC)n microsatellite sequences, and no read pairs spanning the breakpoint between *R2d2* and flanking sequence were identified.

In order to obtain a more precise estimate of the molecular age of the duplication event we assembled *de novo* a total of 16.9 kbp of intergenic and intronic sequence in 8 regions across the *R2d* unit from diverse samples and constructed phylogenetic trees. The trees cover 17 *R2d1* or *R2d2* haplotypes, 13 from inbred strains and 4 from wild mice. The sequence of *Mus caroli* (diploid copy number 2) is used as an outgroup. A concatenated tree is shown in **Figure 1C**. Using 5.0 ± 1.0 million years before present (Mya) as the estimated divergence date for *M. caroli* and *M. musculus* (Suzuki *et al*. 2004; Chevret *et al*. 2005), Bayesian phylogenetic analysis with BEAST v1.8 (Drummond *et al*. 2012) yields 1.6 Mya (95% HPD 0.7 — 5.1 Mya) as the estimated age of the duplication event that gave rise to *R2d1* and *R2d2*. Although the assumption of a uniform molecular clock may not be strictly fulfilled for *R2d1* and *R2d2*, the totality of evidence – from presence/absence data across the mouse phylogeny, paralog-specific TE insertions, and sequence divergence between paralogs – strongly supports the conclusion that R2d was first duplicated within the last 2 My in the common ancestor of *M. musculus* and *M. spretus*.

For clarity, **Figure 1D** illustrates diploid copy number states that will be referenced in the remainder of the manuscript. Hereafter we refer to diploid copy numbers except when discussing inbred strains (which are effectively haploid).

### *R2d* contains the essential gene *Cwc22*

The *R2d* unit encompasses one protein-coding gene, *Cwc22*, which encodes an essential mRNA splicing factor (Yeh *et al*. 2010). The gene is conserved across eukaryotes and is present in a single copy in most non-rodent species represented in the TreeFam database (http://www.treefam.org/family/TF300510 (Li 2006)). Five groups of *Cwc22* paralogs are present in mouse genomes: the copies in *R2d1* (*Cwc22*^*R2d1*^) and *R2d2* (*Cwc22*^*Ridi*^) plus retrotransposed copies in one locus at chr2: 83.9 Mbp and at two loci on the *X* chromosome (**Figure 2A**).

**Figure 2.**
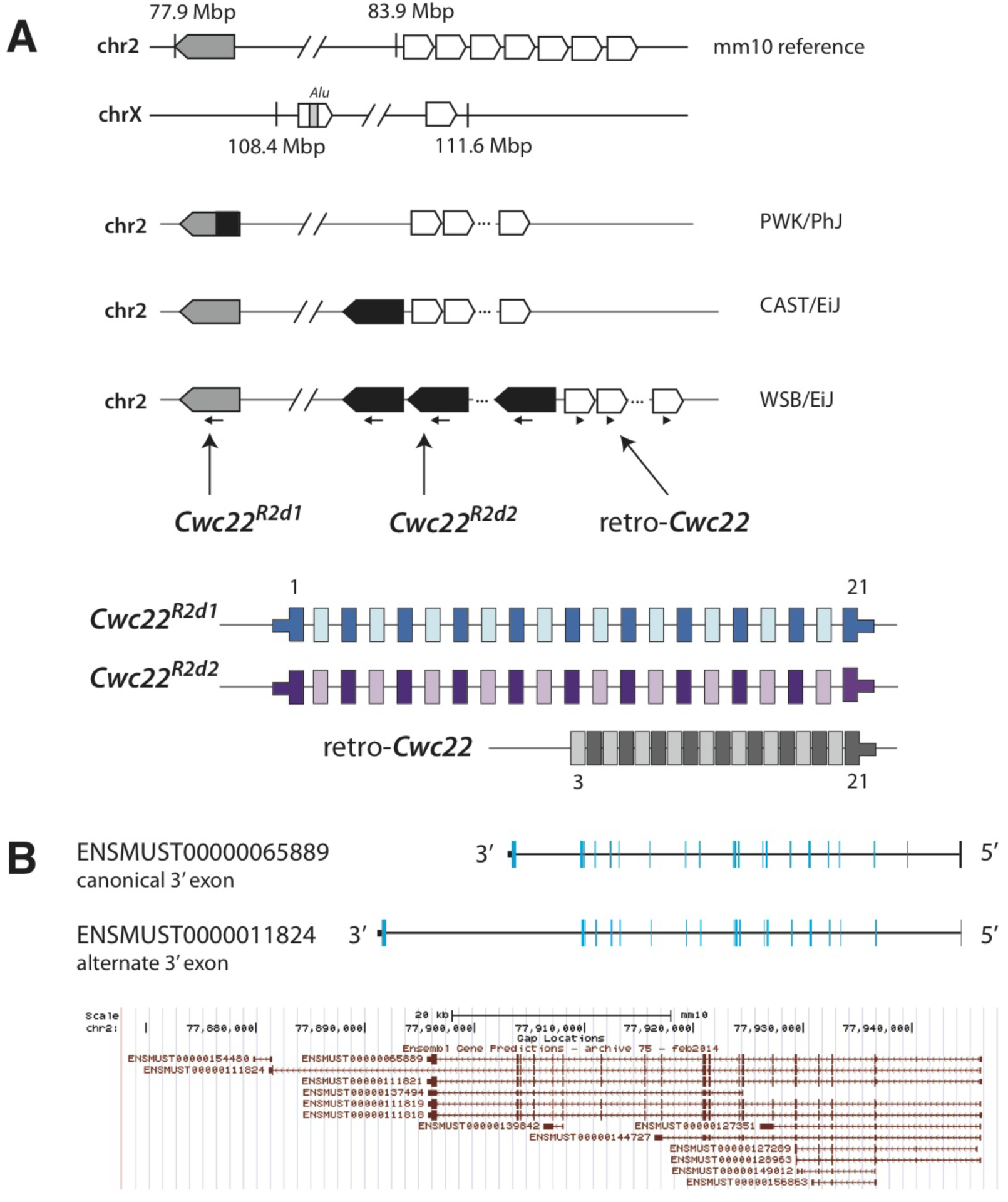
*Cwc22* paralogs in the mouse genome. **(A)** Location and organization of *Cwc22* gene copies present in mouse genomes. The intact coding sequence of *Cwc22* exists in in both *R2d1* (grey shapes) and *R2d2* (black shapes). Retrotransposed copies (empty shapes) exist in two loci on chrX and one locus on chr2, immediately adjacent *R2d2*. Among the retrotransposed copies, coding sequence is intact only in the copy on chr2. Exon numbers are shown in grey above transcript models. **(B)** Alternate transcript forms of *Cwc22*, using different 3’ exons. Coding exons shown in blue and untranslated regions in black. All Ensembl annotated transcripts are shown in the lower panel (from UCSC Genome Browser.)

The three retrogenes are located in regions with no sequence similarity to each other, indicating that each represents an independent retrotransposition event. The copy on chr2 was subsequently expanded by further segmental duplication and now exists (in the reference genome) in 7 copies with >99.9% mutual similarity. The two retrotransposed copies on chrX are substantially diverged from the parent gene (<90% sequence similarity), lack intact open reading frames (ORFs), have minimal evidence of expression among GenBank cDNAs, and are annotated as likely pseudogenes (Pei *et al*. 2012). We therefore restricted our analyses to the remaining three groups of *Cwc22* sequences, all on chr2.

The canonical transcript of *Cwc22*^*R2d1*^ (ENSMUST00000065889) is encoded by 21 exons on the negative strand. The coding sequence begins in the third exon and ends in the terminal exon (**Figure 2B**). Six of the seven protein-coding *Cwc22*^*R2d2*^ transcripts in Ensembl v83 use this terminal exon, while one transcript (ENSMUST0000011824) uses an alternative terminal exon. Alignment of the retrogene sequence (ENSMUST00000178960) to the reference genome demonstrates that the retrogene captures the last 19 exons of the canonical transcript ‒ that is, the 19 exons corresponding to the coding sequence of the parent gene.

### Copy number at *R2d2* is highly polymorphic in *M. musculus*

We previously demonstrated that haploid copy number of *R2d* ranges from 1 in the reference strain C57BL/6J and classical inbred strains A/J, 129S1/SvImJ, NOD/ShiLtJ, NZO/HILtJ; to 2 in the wild-derived strain CAST/EiJ; to 34 in the wild-derived strain WSB/EiJ. Using linkage mapping in two multiparental genetic reference populations, the Collaborative Cross (Collaborative Cross Consortium 2012) and Diversity Outbred (Svenson *et al*. 2012), we showed that, for the two strains with haploid copy number >1, one of the copies maps to *R2d2* while all extra copies map to the *R2d2* locus at chr2: 83 Mbp (Didion *et al*. 2015). *Cwc22* was recently reported to have diploid copy number as high as 83 in wild *M. m. domesticus* (Pezer *et al*. 2015). In whole-genome sequence data from more than sixty mice from both laboratory stocks and natural populations (**Supplementary Table 1**), we have observed zero instances in which the *R2d* copy in *R2d1* is lost. We conclude that diploid copy number >2 indicates at least one copy of *R2d* is present in *R2d1* (**Figure 1D**).

In order to understand the evolutionary dynamics of copy-number variation at *R2d2*, we investigated the relationship between copy number and the local phylogeny in the *R2d2* candidate region. In particular, we sought evidence for or against a single common origin for each of the copy-number states at *R2d2* which are derived with respect to the *M. spretus* - *M. musculus* common ancestor (**Figure 1D**). If a derived copy-number state has a single recent origin, it should be associated with a single haplotype at *R2d2*. If a derived copy-number state arises by recurrent mutation, the same copy number should be associated with multiple haplotype backgrounds and possibly in multiple populations.

The extent of *R2d* copy-number variation in *M. musculus*, as estimated on a continuous scale by qPCR, is shown in **Figure 3A.** (Note that the qPCR readout is proportional to copy number on the log scale. Extrapolation to integer copy number is imprecise for copy numbers greater than ~6.) We confirmed that *R2d2* maps to chr2: 83 Mbp by performing association mapping between SNP genotypes from the MegaMUGA array (Morgan *et al*. 2016) and the qPCR readout (**Figure 3B**).

**Figure 3.**
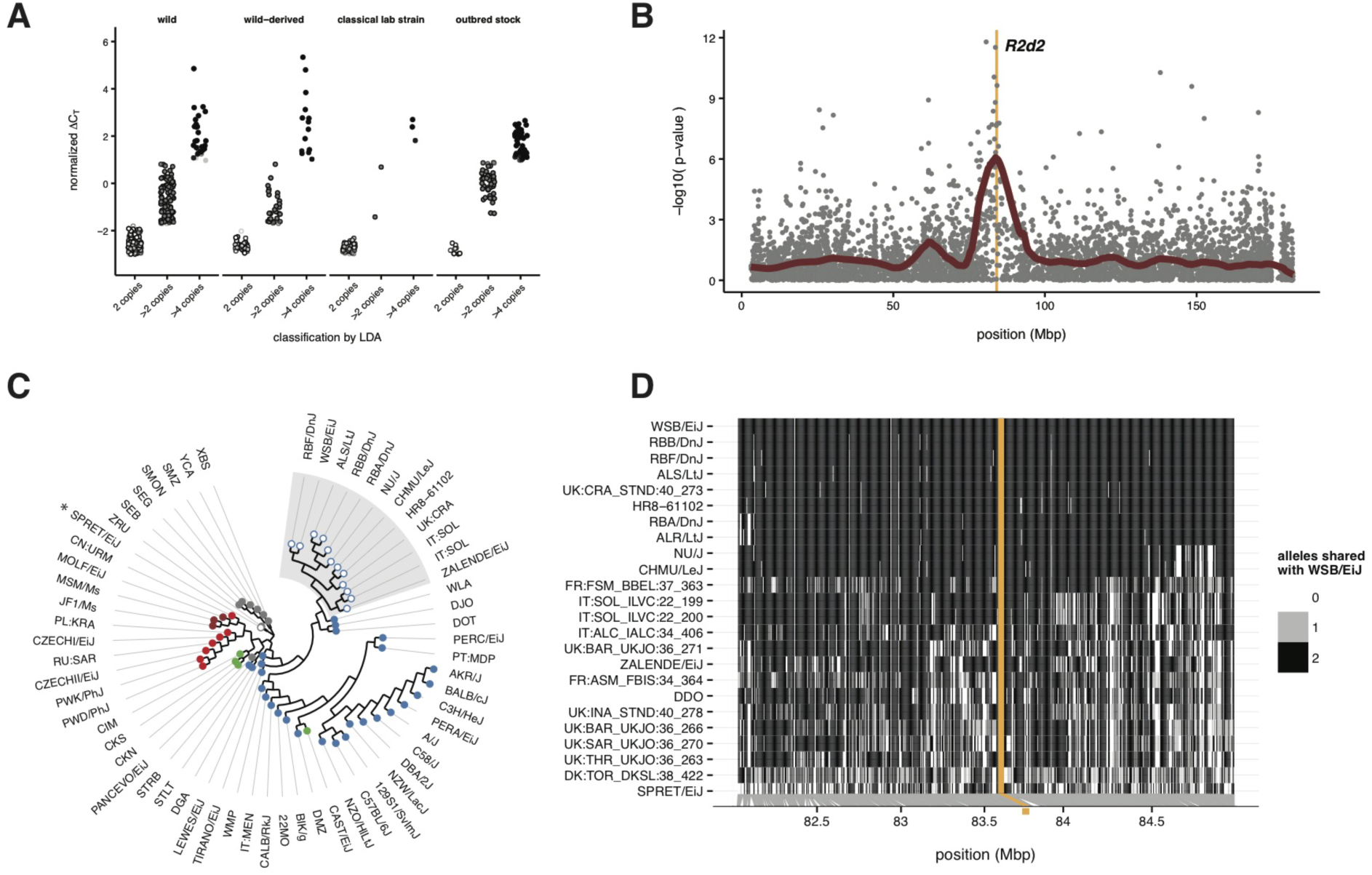
Copy-number variation of *R2d* in mouse populations worldwide. **(A)** Copy-number variation as measured by quantitative PCR. The normalized deltaCt value is proportional to log2(copy number). Samples are classified as having 2 diploid copies, >2 copies or >4 copies of *R2d* using linear discriminant analysis (LDA). **(B)** Fine-mapping the location of *R2d2* in 83 samples genotyped on the Mouse Diversity Array (MDA). Grey points give nominal p-values for association between *R2d* copy number and genotype; red points show a smoothed fit through the underlying points. The candidate interval for *R2d2* from Didion *et al*. (2015), shown as an orange shaded box, coincides with the association peak. **(C)** Local phylogeny at chr2: 83-84 Mbp in 62 wild-caught mice and laboratory strains. Tips are colored by subspecies of origin: *M. m. domesticus*, blue; *M. m. musculus*, red; *M. m. castaneus*, green; *M. m. molossinus*, maroon; outgroup taxa, grey. Individuals with >4 diploid copies of *R2d* are shown as open circles. **(D)** Haplotypes of laboratory strains and wild mice sharing a high-copy allele at *R2d2*. All samples share a haplotype over the region shaded in orange.

To test the hypothesis that losses of *R2d2* (diploid copy number < 4, at least one chromosome with zero copies in *R2d2*, Figure 1D) have a single origin, we examined their distribution across the three well-differentiated subspecies of *M. musculus*. Losses of *R2d2* occur in all subspecies of *M. musculus*, in populations that span its geographic range (**Supplementary Table 3**). Based on this distribution and a previous observation that no common haplotype is shared in samples with low copy number in *M. m. domesticus* (Didion *et al*. 2016), we reject the hypothesis of single origin and conclude that *R2d2* has been lost multiple times on independent lineages in each subspecies.

We performed a similar analysis to test the hypothesis that *R2d2* alleles with high copy number (diploid copy number >4, **Figure 1D**; hereafter *“R2d2*^*HC*^”) have a single origin. First we observed that *R2d2*^*HC*^ alleles are confined with few exceptions to *M. m. domesticus* (**Supplementary Table 3**). The best-associated SNP on the MegaMUGA array (JAX00494952) only weakly tags copy number (*r*^2^ = 0.137), but ascertainment bias on the MUGA platform (Morgan *et al*. 2016) makes local LD patterns difficult to interpret. To examine further, we constructed a neighbor-joining phylogenetic tree for the region containing *R2d2* (chr2: 83 - 84 Mb) using genotypes from the 600K-SNP Mouse Diversity Array (Yang *et al*. 2011). We restricted our attention to inbred strains or wild mice with homozygous, non-recombinant haplotypes in the target region. Twelve samples with *R2d2*^*HC*^ alleles, both wild mice and laboratory stocks, cluster in a single clade (**Figure 3C**). (A single *M. spretus* strain, SPRET/EiJ, also carries an *R2d2*^*HC*^ allele, but see **Discussion**).

Next we expanded the analysis to include an additional 11 samples with *R2d2*^*HC*^ alleles and evidence of heterozygosity around *R2d2*. The total set of 24 samples includes 7 wild-derived laboratory strains (DDO, RBA/DnJ, RBB/DnJ, RBF/DnJ, WSB/EiJ, ZALENDE/EiJ and SPRET/EiJ), 4 classical inbred strains (ALS/LtJ, ALR/LtJ, CHMU/LeJ and NU/J), a line derived from the ICR:HsD outbred stock (HR8; Swallow *et al*. 1998) and 12 wild-caught mice. All 24 samples with *R2d2*^*HC*^ alleles share an identical haplotype across a single 21 kbp interval, chr2: 83,896,447 - 83,917,565 (GRCm38/mm10 coordinates) (**Figure 3D**). These analyses support a single origin for *R2d2*^*HC*^ alleles within *M. m. domesticus*.

### *Cwc22* is intact in and expressed from all *R2d* paralogs, and fast-evolving in rodents

To identify the coding sequence of *Cwc22*^*R2d2*^ we first aligned the annotated transcript sequences of *Cwc22*^*R2d1*^ from Ensembl to our *R2d2* contig. All 21 exons present in *R2d1* are present in *R2d2*. We created a multiple sequence alignment and phylogenetic tree of *Cwc22* cDNAs and predicted amino acid sequences from *Cwc22*^*R2d1*^*, Cωcll*^*R2d2*^, retro-Cwc22, and *Cwc22* orthologs in 19 other placental mammals, plus opossum, platypus and finally chicken as an outgroup (**Supplementary Figure 3**). An open reading frame (ORF) is maintained in all three *Cwc22* loci in mouse, including the retrogene. Information content of each column along the alignment (**Supplementary Figure 4**) reveals that sequence is most conserved in two predicted conserved domains, MIF4G and MA3, required for *Cwc22s* function in mRNA processing (Yeh *et al*. 2010).

Next we examined public RNA-seq data from adult brain and testis in inbred strains with one or more copies of *R2d2* for evidence of transcription of each *Cwc22* family member. We identified several novel transcript isoforms specific to *R2d2* arising from two intron-retention events and one novel 3’ exon (**Figure 4A**). The 18^th^ intron is frequently retained in *Cwc22*^*R2d2*^ transcripts, most likely due to an A>G mutation at the 5’ splice donor site of exon 17 in *Cwc22*^*R2d2*^. The 12^th^ intron is also frequently retained. While we could not identify any splice-region variants near this intron, it contains an ERV insertion that may interfere with splicing (**Figure 4A**). Both intron-retention events would create an early stop codon. Finally, we find evidence for a novel 3’ exon that extends to the boundary of the *R2d* unit and is used exclusively by *Cwc22*^*R2d2*^ (**Figure 4A**).

**Figure 4.**
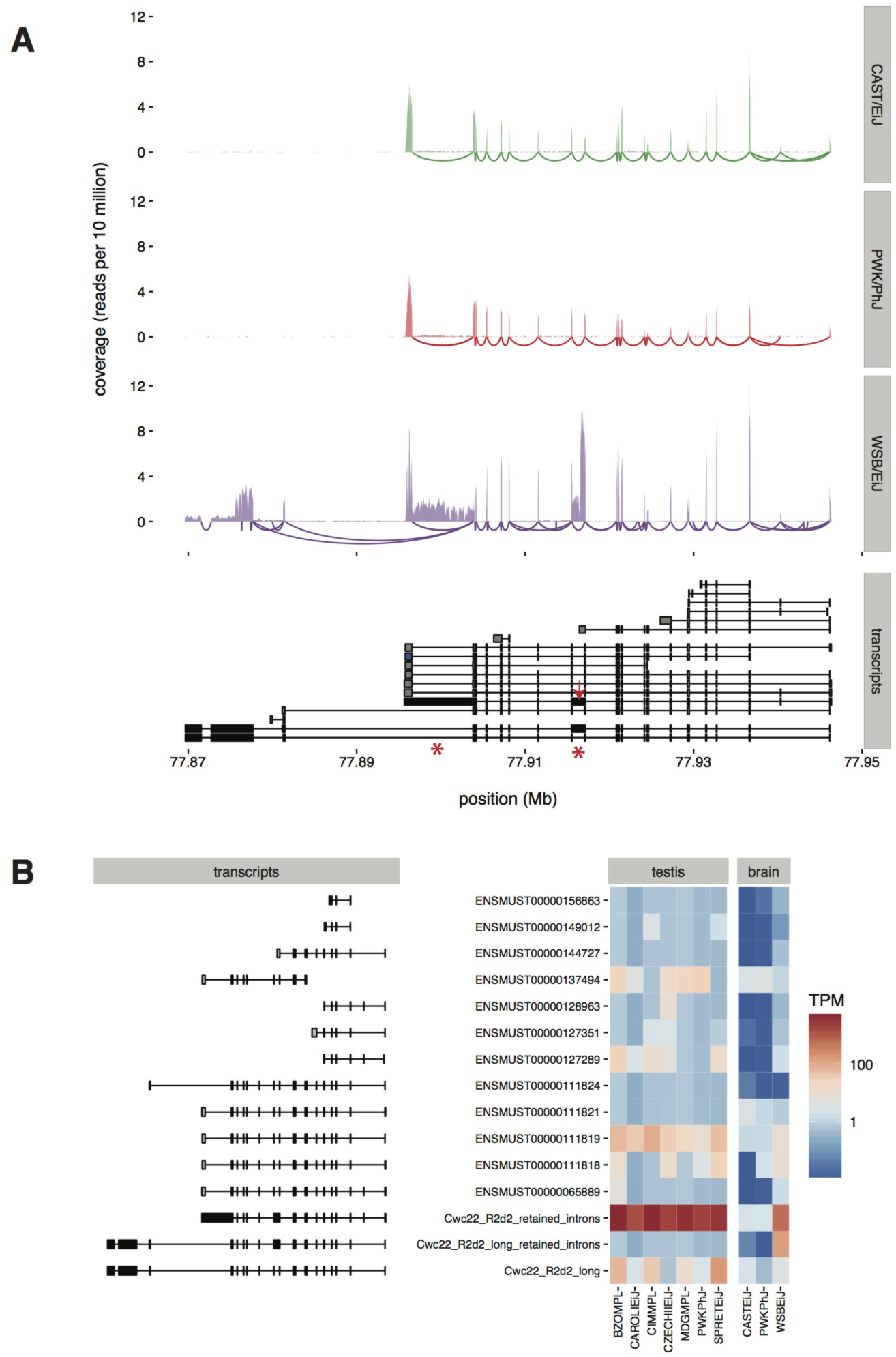
Expression of *Cwc22* isoforms. **(A)** Read coverage and splicing patterns in *Cwc22* in adult mouse brain from three wild-derived inbred strains. Swoops below x-axis indicate splicing events supported by 5 or more split-read alignments. Known transcripts of *Cwc22*^*R2d1*^ (grey, from Ensembl), inferred transcripts from *Cwc22*^*R2d2*^ (black) and the sequence of retro-*Cwc22* mapped back to the parent gene (blue) are shown in the lower panel. Red stars indicate retained introns; red arrow indicates insertion site of an ERV in *R2d2*. **(B)** Estimated relative expression of *Cwc22* isoforms (y-axis) in adult mouse brain and testis in wild-derived inbred strains (x-axis). TPM, transcripts per million, on log10 scale.?

We estimated the expression of the various isoforms of *Cwc22*^*R2d1*^, *Cwc22*^*R2d2*^ and retro-*Cwc22* in adult brain and testis. For brain we obtained reads from 8 replicates (representing both sexes) on 3 inbred strains, and for testis a single replicate on 23 inbred strains and estimated transcript abundance using the kallisto package (Bray *et al*. 2015). Briefly, kallisto uses an expectation-maximization (EM) algorithm to accurately estimate the abundance of a set of transcripts by distributing the “weight” of each read across all isoforms with whose sequence it is compatible. *Cwc22* is clearly expressed from all three paralogs in both brain and testis (**Figure 4B**). However, both the total expression and the pattern of isoform usage differ by tissue and copy number.

Maintenance of an ORF in all *Cwc22* paralogs for >2 My is evidence of negative selection against disrupting mutations in the coding sequence, but long branches within the rodent clade in **Supplementary Figure 3** suggest that *Cwc22* may also be under relaxed purifying selection or positive selection in rodents. The rate of evolution of *Cwc22* sequences in mouse is faster than in the rest of the tree (*χ*^2^ = 4.33, df = 1, *p* = 0.037 by likelihood ratio test).

### Phylogenetic discordance in *R2d1* is due to non-allelic gene conversion

The topology of trees across *R2d* is generally consistent: a long branch separating the single *M. caroli* sequence from the *M. musculus* sequences, and two clades corresponding to *R2d1*- and *R2d2*-like sequences. However, we observed that the affinities of some *R2d* paralogs change along the sequence (**Figure 5A**), a signature of non-allelic (*i.e*. inter-locus) gene conversion. In this context, we use “gene conversion” to describe a non-reciprocal “copy-and-paste” transfer of sequence from one donor locus into a different, homologous receptor locus, without reference to a specific molecular mechanism (Chen *et al*. 2007).

**Figure 5.**
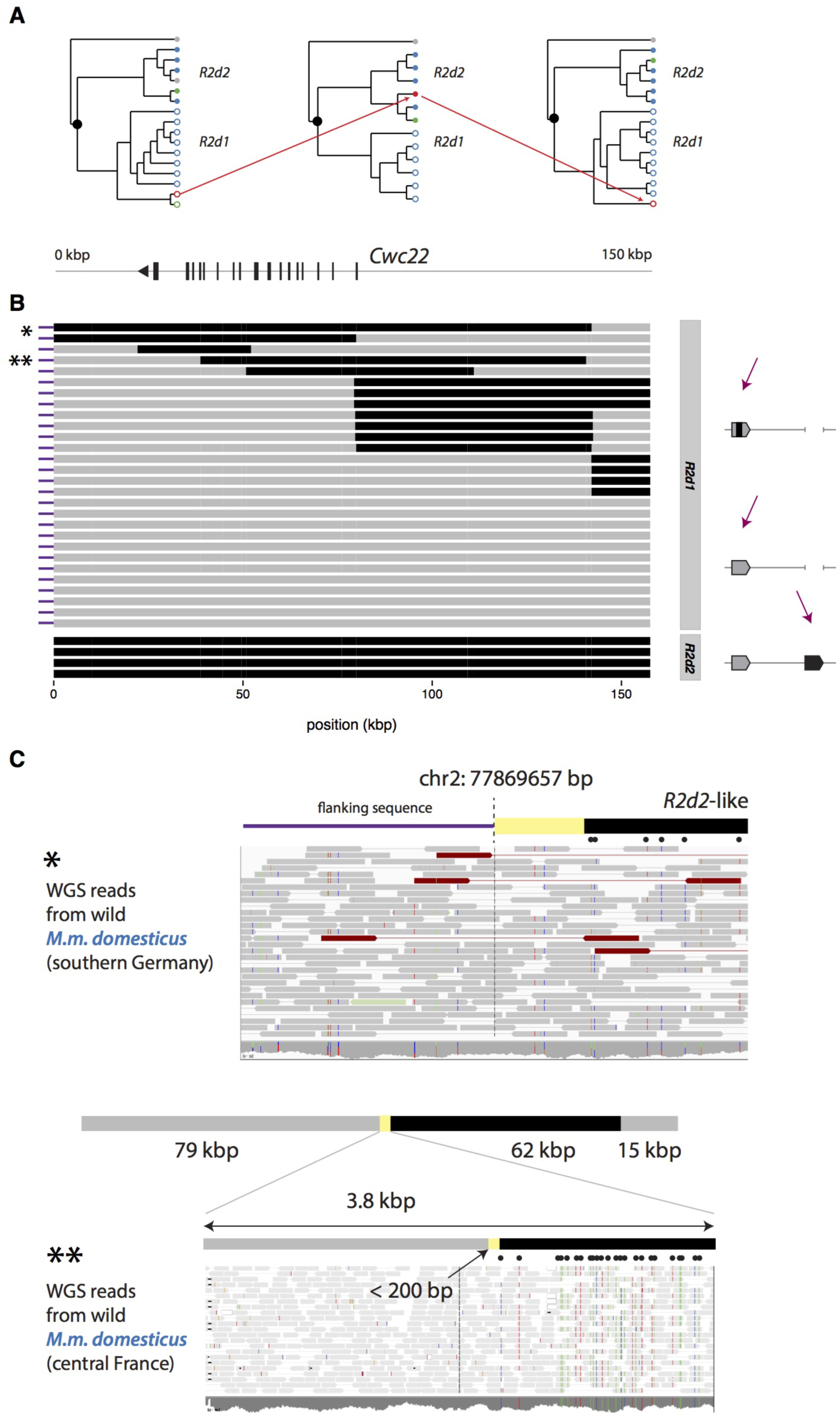
Signatures of non-allelic gene conversion between *R2d1* and *R2d2*. **(A)** Phylogenetic trees for three representative intervals across *R2d*. Sequences are labeled according to their subspecies of origin using the same color scheme as in Figure 1; open circles are *R2d1*-like sequences and closed circles are *R2d2*-like. Trees are drawn so that *M. caroli*, the outgroup species used to root the trees, is always positioned at the top. The changing affinities of PWK/PhJ (red) and CAST/EiJ (green) along *R2d* are evidence of non-allelic gene conversion. **(B)** *R2d* sequences from 20 wild-caught mice and 5 laboratory inbred strains. Each track represents a single chromosome; grey regions are classified as *R2d1*-like based on manual inspection of sequence variants, and black-regions *R2d2*-like. Upper panel shows sequences from samples with a single copy of *R2d*, residing in *R2d1*. Lower panel shows representative *R2d2* sequences for comparison. Asterisks indicate samples for which read alignments are shown in panel C. **(C)** Upper panel, paire-end read alignments (visualized with IGV) across the proximal boundary (dashed line) of *R2d1* in a sample with a conversion tract extending to the boundary. Positions of derived variants shared with *R2d2* are indicated by black dots. Lower panel, read alignments across the boundary of a non-allelic gene conversion tract. *R2d1* sequence from a single chromosome is a mosaic of *R2d1*-like (grey) and Rldi-like (black) segments. A magnified view of read pairs in the 3.8 kbp surrounding the proximal boundary of the tract shows read pairs spanning the junction. Black dots indicate the position of derived alleles diagnostic for *R2d2*. The precise breakpoint lies somewhere in the yellow shaded region between the last *R2d1*-specific variant and the first R2d2-specific variant.

To investigate further, we inspected patterns of sequence variation in whole-genome sequencing data from 15 wild-caught mice, 2 wild-derived inbred strains, and 22 classical inbred strains of mice with diploid *R2d* copy number 2. We first defined 1,411 pairwise single-nucleotide differences between *R2d2* and *R2d1* for which *R2d2* has the derived allele with respect to *M. caroli*. Then we tested for the presence of the derived allele, ancestral allele or both at each site in each sample. Finally we identified conversion tracts by manual inspection as clusters of derived variants shared with *R2d2* (**Supplementary Figure 5**).

This analysis revealed non-allelic gene conversion tracts on at least 9 chromosomes (Figure 5B). The conversion tracts range in size from approximately 1.2 kbp to 119 kbp. The boundaries of several tracts are shared within populations, suggesting that they are identical by descent. We excluded the possibility of complementary losses from *R2d1* and *R2d2* - which would leave similar patterns of sequence variation - by finding read pairs spanning the boundary between *R2d1* and flanking sequence, and between *R2d1*- like and *R2d2*-like tracts on the same chromosome (examples shown in **Figure 5C**).

The conversion tracts we detected are orders of magnitude longer than the 15 to 750 bp reported in recent studies of allelic gene conversion at recombination hotspots in mouse meiosis (Cole *et al*. 2010, 2014). We require the presence of *R2d2*-diagnostic alleles at two or more consecutive variants to declare a conversion event, and these variants occur at a rate of approximately 1 per 100 bp, so the smallest conversion tracts we could theoretically detect are on the order of 200 bp in size. Even if we require only a single variant to define a conversion tract, all samples without a long conversion tract share fewer than 55 and most fewer than 10 (of 1,411) derived alleles with *R2d2*, of which all are also shared by multiple other samples from different populations (**Supplementary Figure 5**). This pattern indicates that those sites in fact represent either artifacts (from mis-assignment of ancestral and derived alleles) or homoplasy rather than short gene conversions.

Four conversion tracts partially overlap the *Cwc22* gene to create a sequence that is a mosaic of *R2d1*- and *R2d2*-like exons (**Figure 5B**). Recovery of *Cwc22* mRNA in an inbred strain with a mosaic sequence (PWK/PhJ, see section *“Cwc22* is intact and expressed”) indicates that its exons are intact and properly oriented in *cis*. The presence of both *R2d1*- and *R2d2*-like sequence in extant *M. musculus* lineages with a 2 diploid copies of *R2d* further reinforces our conclusion that the duplication is indeed ancestral to the divergence of *M. musculus*.

In addition to exchanges between *R2d1* and *R2d2*, we identified an instance of exchange between *R2d2* and the adjacent retrotransposed copy of *Cwc22* in a single *M. m. domesticus* individual from Iran (IR:AHZ_STND:015; Supplementary Figure 6). This individual carries a rearrangement that has inserted a 30 kbp fragment corresponding to the 3’ half of *Cwc22*^*R2d2*^ into the retro-*Cwc22* locus, apparently mediated by homology between the exons of *Cwc22*^*R2d2*^ and retro-*Cwc22*.

**Figure 6.**
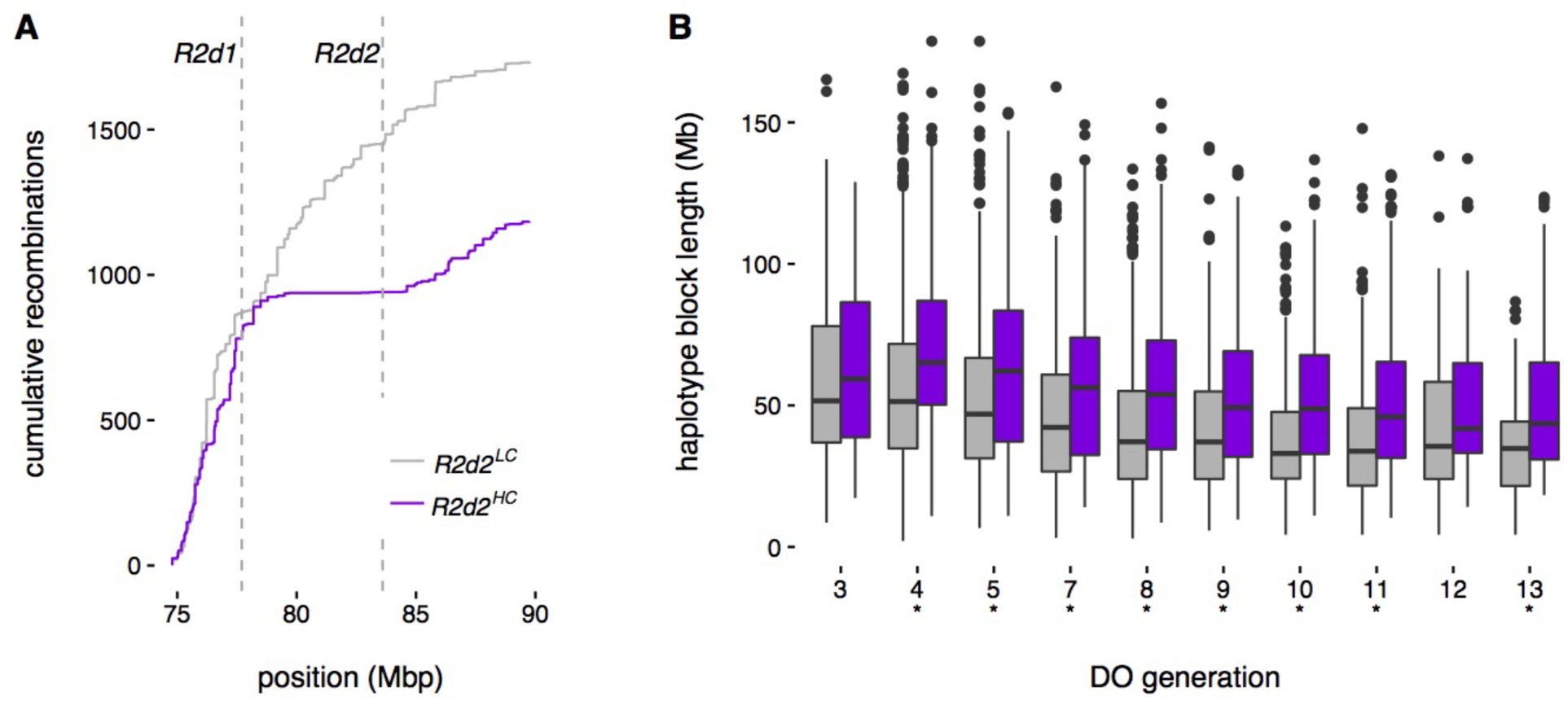
Suppression of crossing-over around *R2d2*. **(A)** Cumulative number of unique recombination events in the middle region of chr2 in genomes of 4,640 Diversity Outbred mice. Recombination events involving the high-copy-number WSB/EiJ haplotype are shown in purple and all other events in grey. Dashed vertical lines indicate the position of *R2d1* (left) and *R2d2* (right). **(B)** Distribution of haplotype block sizes at *R2d2* in selected generations of the DO, for *R2d2*^*HC*^ (WSB/EiJ, purple) versus *R2d2*^*LC*^ (the other seven founder haplotypes, grey). Asterisks indicate generations in which the length distributions are significantly different by Wilcoxon rank-sum test.

Based on a previous observation that the rate of meiotic recombination is reduced near clusters of segmental duplications (Liu *et al*. 2014), we tested whether the region around *R2d2* has lower recombination when an *R2d2*^*HC*^ allele is present. Understanding patterns of recombination at *R2d2* is important for interpreting levels of sequence and haplotype diversity in the surrounding region.

First we analyzed local recombination rate in the Diversity Outbred (DO) population. The DO is an outbred stock derived from eight inbred founder strains (including one, WSB/EiJ, with an *R2d2*^*HC*^ allele) and maintained by random mating with 175 breeding pairs; at each generation, one male and one female offspring are chosen from each mating and randomly paired with a non-sibling to produce the next generation (Svenson *et al*. 2012). **Figure 6A** shows the cumulative distribution of 2,917 recombination events on central chromosome 2, stratified according to *R2d2* copy number of the participating haplotypes. The recombination map has a pronounced plateau in the region between *R2d1* and approximately 1 Mb distal to *R2d2* (dashed lines) for *R2d2*^*HC*^ haplotypes, but not *R2d2*^*LC*^ haplotypes. As a result, *R2d2*^*HC*^ haplotype blocks overlapping *R2d2* are significantly shorter than *R2d2*^*LC*^ haplotype blocks (*p* < 0.01 by Wilcoxon rank-sum tests with Bonferroni correction) in 8 of the 10 generations sampled (**Figure 6B**). The difference arose early in the breeding of the DO and persists through the most recent generation for the randomized breeding scheme was maintained (FPMdV, unpublished).

Second we re-examined genotype data from 11 published crosses in which at least one parent was segregating for an *R2d2*^*HC*^ allele. Whereas in the DO we used haplotype block length as a proxy for recombination rate, in these F2 and backcross designs we can directly estimate the recombination fraction across *R2d2* and compare it to its expected value in the absence of an *R2d2*^*HC*^ allele (**Supplementary Figure 7**). In 9 of 11 crosses examined, the observed recombination fraction is lower than the expected (*p* < 0.032, one-sided binomial test).

### The genomic region containing *R2d2* is structurally unstable but has low sequence diversity

The extent of copy-number polymorphism involving *R2d2* suggests that it is intrinsically unstable. Consistent with these observations, we find that the rate of *de novo* copy-number changes at *R2d2* is extremely high in laboratory populations (**Figure 7**). In 183 mice sampled from the DO population we identified and confirmed through segregation analysis 8 new alleles, each with distinct copy number and each occurring in an unrelated haplotype (**Supplementary Table 4**). Without complete pedigrees and genetic material from breeders a direct estimate of the mutation rate in the DO is not straightforward to obtain. However, since the population size is known, we can make an analogy to microsatellite loci (Moran 1975) and estimate the mutation rate via the variance in allele sizes: 3.2 mutations per 100 transmissions (3.2%) (95% bootstrap CI 1.1% — 6.0%).

**Figure 7.**
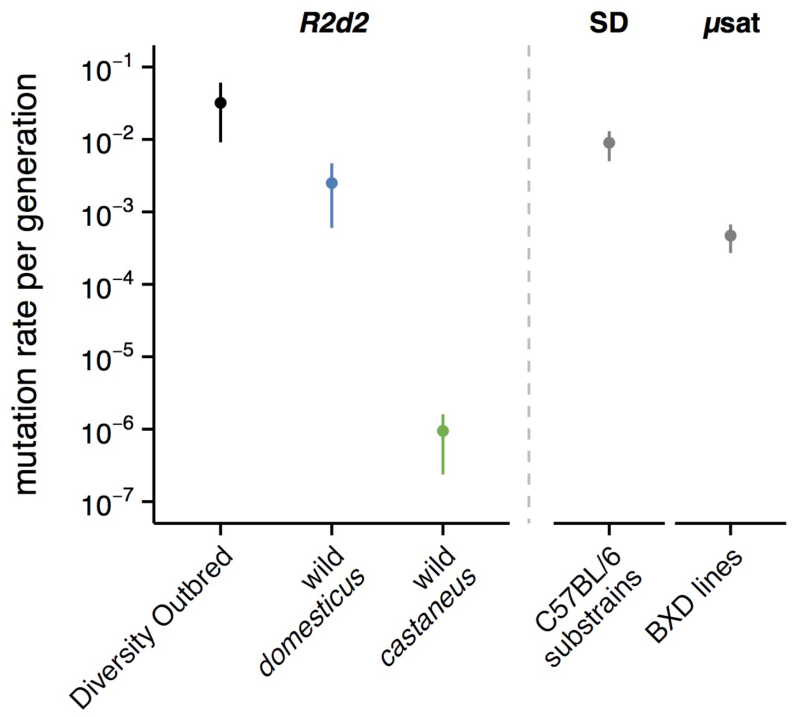
ate of *de novo* copy-number changes at *R2d2*. Estimâtes of per-generation mutation rate for CNVs at *R2d2* (±1 bootstrap SE) in the Diversity Outbred population; among wild *M. m. domesticus;* and among wild *M. m. castaneus*. For comparison, mutation rates are shown for the CNV with the highest rate of recurrence in a C57BL/6J pedigree (Egan *et al*. 2007) and for a microsatellite whose mutation rate was estimated in the BXD panel (Dallas 1992).

Structural instability in this region of chromosome 2 extends outside the *R2d2* locus itself. Less than 200 kbp distal to *R2d2* is another segmental duplication (**Figure 8B**, grey shaded region) — containing a retrotransposed copy of *Cwc22* — that is present in 7 tandem copies in the reference genome. That region, plus a further 80 kbp immediately distal to it, is copy-number polymorphic in wild *M. m. domesticus* and wild *M. m. castaneus* (**Figure 8B**). Instability of the region over longer timescale is demonstrated by the disruption, just distal to the aforementioned segmental duplication, of a syntenic block conserved across all other mammals (**Supplementary Figure 1**).

**Figure 8.**
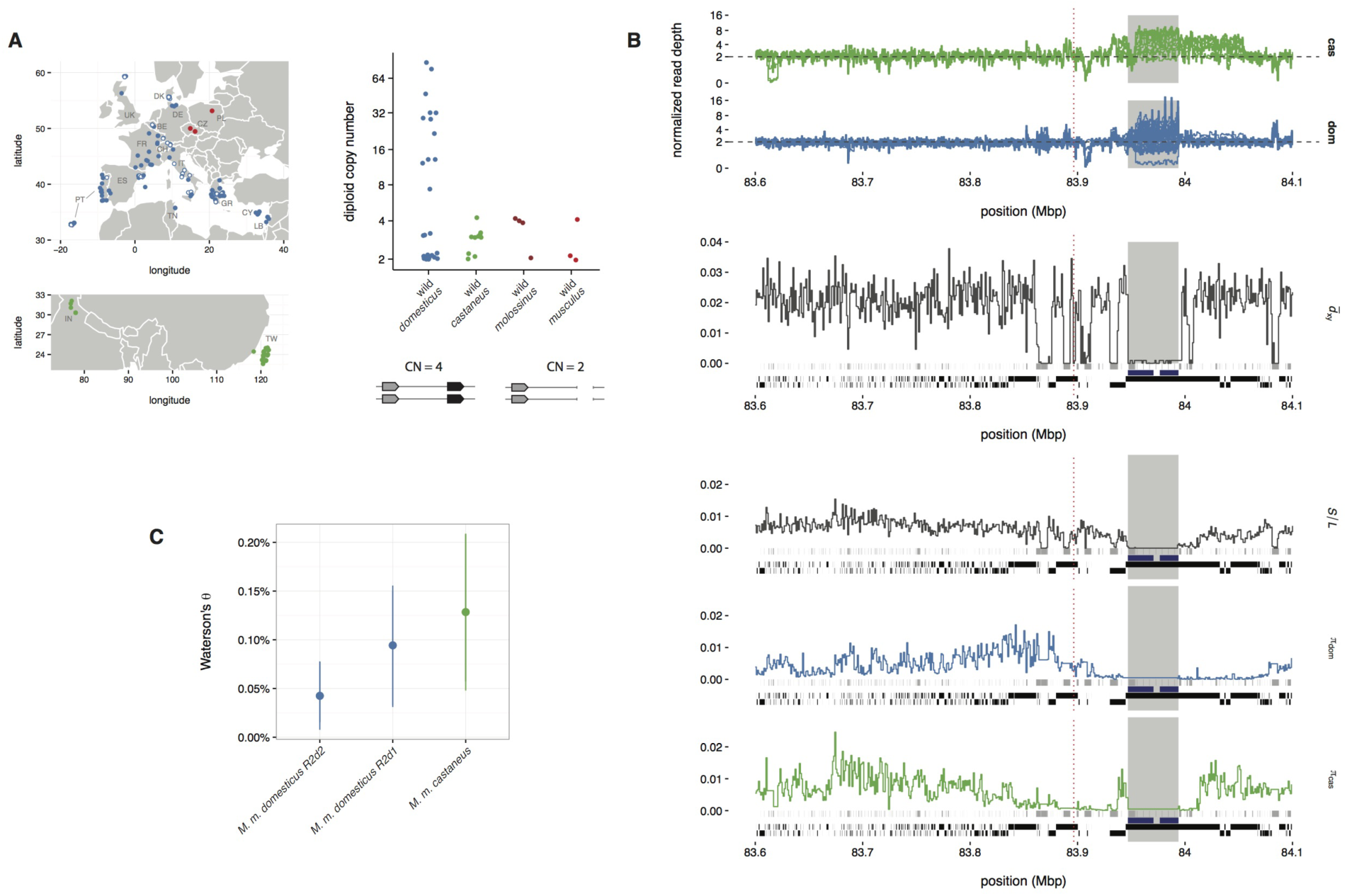
Sequence and structural diversity around *R2d2*. **(A)** Geographic origin of wild mice used in this study, color-coded by subspecies (blue, *M. m. domesticus*; red, *M. m. musculus*; green, *M. m. castaneus*). Diploid copy number of the *R2d* unit is shown for wild samples for which integer copy-number estimâtes are available: 26 *M. m. domesticus* and 10 *M. m. castaneus* with whole-genome sequencing data, and representatives from *M. m. molossinus* and *M. m. musculus* for comparison. Schematic shows the *R2d1/R2d2* configurations corresponding to diploid copy numbers of 2 and 4. **(B)** Profiles of read depth (first two panels), average sequence divergence to outgroup species *M. caroli* (*d*_xy_, third panel), number of segregating sites per base (*S/L*, fourth panel) and within-population average heterozygosity (*π*, fifth and sixth panels). The region shown is 500 kbp in size; the insertion site of *R2d2* is indicated by the red dashed line. Grey boxes along baseline show positions of repetitive elements (from UCSC RepeatMasker track); black boxes show non-recombining haplotype blocks. Blue bars indicate the position of 7 tandem duplications in the mm10 reference sequence with >99% mutual identity, each containing a copy of retro-*Cwc22*. Grey shaded region indicates duplicate sequence absent from *M. caroli*. **(C)** Estimated per-site nucleotide diversity within *M. m. domesticus *R2d1*, M. m. domesticus R2d2* and *M. m. castaneus R2d2*.

Despite the high mutation rate for structural variants involving *R2d2* and nearby sequences, sequence diversity at the nucleotide level is modestly reduced relative to diversity in *R2d1* and relative to the genome-wide average in *M. m. domesticus*. In a 200 kbp region containing the *R2d2* insertion site at its proximal end, *π* (an estimator of average heterozygosity) in *M. m. domesticus* reduced from approximately 0.3% (comparable to previous reports in this subspecies, (Salcedo *et al*. 2007)) to nearly zero (**Figure 8B**). Divergence between *M. musculus* and *M. caroli* is similar to its genome-wide average of ~ 2.5% over the same region.

Estimation of diversity *within* a duplicated sequence such as *R2d* is complicated by the difficulty of distinguishing allelic from paralogous variation. To circumvent this problem we split our sample of 26 wild *M. m. domesticus* into two groups: those having *R2d1* sequences only, and those having both *R2d1* and *R2d2* sequences. Within each group we counted the number of segregating sites among all *R2d2* copies, using nearby fixed differences between *R2d1* and *R2d2* to phase sites to *R2d2* (see Methods for details), and used Watterson’s estimator to calculate nucleotide diversity per site. Among *R2d1* sequences, *θ* = 0.09% ± 0.03% versus *θ* = 0.04% ± 0.02% among *R2d2* sequences (Figure 8C) and *Θ =* 0.13% ± 0.04% among *R2d2* sequences in *M. m. castaneus*.

### The “revolving door” affects at least 0.8% of the mouse genome

The superposition of duplication, non-allelic gene conversion, and loss — dubbed the “genomic revolving door” (Demuth *et al*. 2006) — can obscure the true history of multicopy sequences. Over short evolutionary times, duplication and loss may be further confounded with incomplete lineage sorting, or the stochasitic distribution of ancestral polymorphisms along descendant lineages (Pamilo and Nei 1988).

Most previous studies of gene gain and loss have focused on between-species comparisons using finished genome assemblies (Demuth *et al*. 2006; Bailey and Eichler 2006). We sought evidence for the “revolving door” effect over the last ~1 My of evolution within *M. musculus*. To do so we identified regions where the divergence between sequenced samples and the reference genome assembly substantially exceeds what is predicted by those samples' ancestry. We developed a simple and unbiased method to estimate divergence from short sequencing reads without alignment. Briefly, we estimated the proportion of short subsequences (*k*-mers, with *k* = 31) from windows along the reference assembly for which no evidence exists in the reads generated for a sample. This quantity can be rescaled to approximate the divergence between the reference sequence and the template sequence from which reads were generated (see **Methods**). Applied genome-wide to 31 kbp (= 1000×*k*) windows, this method captures the distribution of sequence divergence between the reference assembly (chiefly *M. m. domesticus* in origin) in representative samples from the three subspecies of *M. musculus* and the outgroups *M. spretus* and *M. caroli* (**Figure 9A**).

**Figure 9.**
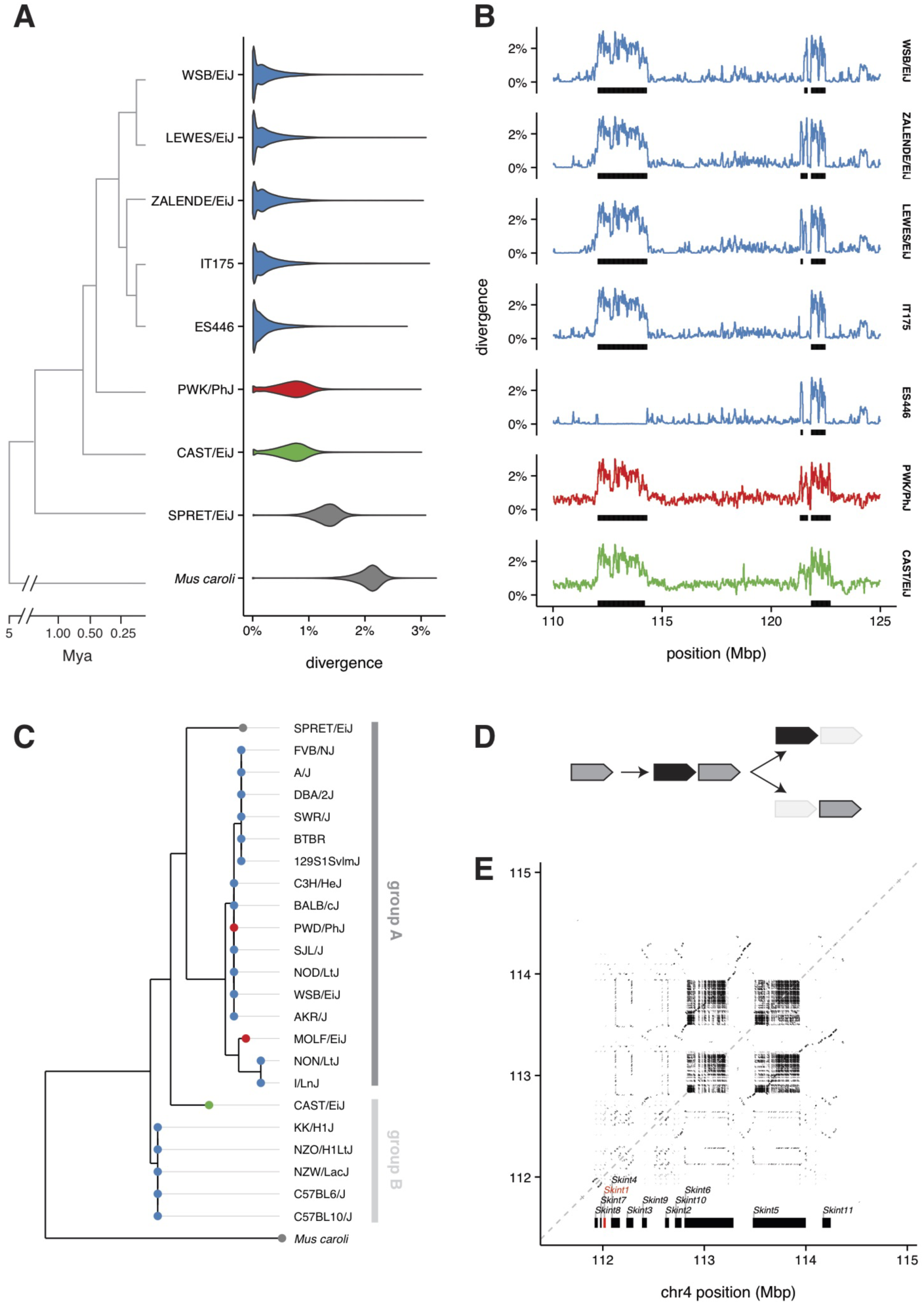
A region of excess sequence divergence on chromosome 4. **(A)** Genome-wide sequence divergence estimates for representative samples from the sub-genus *Mus*. **(B)** Estimated sequence divergence in 31 kbp windows across distal chr4 for the samples in panel A. Divergent regions identified by the hidden Markov model (HMM) are indicated with black bars along the horizontal axis. **(C)** Phylogenetic tree constructed from *Skint1* coding sequences reported in Boyden *et al*. (2008) **(D)** Schematic representation of the process of gene duplication, followed by differential loss of paralogs along independent lineages. **(E)** Dotplot of self-alignment of sequence from the region of distal chr4 containing the *Skint* gene family. Positions of *Skint* genes are indicated along the horizontal axis; *Skint1* highlighted in red.

Divergent regions were identified by fitting a hidden Markov model (HMM) to the windowed divergence profiles for 7 wild or wild-derived samples with available whole-genome sequence (**Figure 9B** and **Supplementary Table 6**). This analysis revealed a striking pattern: over most of the genome, the divergence estimates hover around the genome-wide expectation, but high-divergence windows are clustered in regions 100 kbp to 5 Mbp in size. Our method does not capture signal from structural variation (besides deletions, see **Methods**), and so probably underestimates the true level of sequence divergence. In any case the union of these divergent regions across all seven *M. musculus* samples analyzed covers 0.82% of the reference genome (mean 0.58% per sample). Not surprisingly, divergent regions are enriched for segmental duplications: 39.0% of sequence in divergent regions is comprised of segmental duplications versus a median of 3.4% (central 99% interval 0.2% — 16.7%) in random regions of equal size (*p* < 0.001). Divergent regions also have significant overlap with previously-identified (Liu *et al*. 2014) regions of low recombination (*p* < 0.001). Yet divergent regions are more gene-dense than the genomic background: they contain 3.6×10^−2^ genes/kbp relative to a median 1.6×10^−2^ genes/kbp (central 99% interval 1.1×10^−2^ — 2.7×10^−2^ genes/kbp) genome-wide (*p* = < 0.001). Divergent regions are strongly enriched for genes related to odorant and pheromone sensing (*p* = 1.3×10^−8^) and adaptive immunity (*p* = 3.1×10^−3^).

As a representative example we focus on a divergent region at chr4: 110-115 Mbp (**Figure 9B**). This region contains the 11 members of the *Skint* family of T-cell-borne antigen receptors. The first member to be described, *Skint1*, functions in negative selection in the thymus of T-cells destined for the epidermis (Boyden *et al*. 2008). Coding sequence from 23 inbred strains (including CAST/EiJ, WSB/EiJ, PWD/PhJ and SPRET/EiJ) was reported in Boyden *et al*. (2008). A phylogenetic tree constructed from those sequences, plus *M. caroli* and rat (not shown) as outgroups, reveals the expected pattern of “deep coalescence” (**Figure 9C**): the *M. m. domesticus* sequences are paraphyletic, and some (group A) are more similar to SPRET/EiJ (*M. spretus*) than to their subspecific congeners (group B). Although the level of sequence divergence in the *Skint1* coding sequence was attributed to ancestral polymorphism maintained by balancing selection in the original report, selection would not be expected to maintain diversity in both coding and non-coding sequence equally as we observe in **Figure 9B**. The best explanation for the observed pattern of diversity at *Skint1* is therefore that groups A and B represent “pseudo-orthologs” (Koonin 2005) descended from an ancestral duplication followed by subsequent deletion of different paralogs along different lineages (**Figure 9D**). This conclusion is supported by the structure of the *Skint* region in the mouse reference genome assembly, which reflects the superposition of many duplications and rearrangements (**Figure 9E**).

## DISCUSSION

In this manuscript we have reconstructed in detail the evolution of a multi-megabase segmental duplication (SD) in mouse, *R2d2*. Our findings illustrate the challenges involved in accurately interpreting patterns of polymorphism and divergence within duplicated sequence.

SDs are among the most dynamic loci in mammalian genomes. They are foci for copy-number variation in populations, but the sequences of individual duplicates beyond those present in the reference genome are often poorly resolved. Obtaining the sequence of this “missing genome,” as we have done for *R2d2*, is an important prerequisite to understanding the evolution of duplicated loci. Since each paralog follows a partially independent evolutionary trajectory, individuals in a population may vary both quantitatively (in the number of copies) and qualitatively (in which copies are retained). Cycles of duplication and loss may furthermore lead to the fixation of different paralogs along different lineages. This “genomic revolving door” leaves a signature of polymorphism far in excess of the genome-wide background, due to coalescence between alleles originating from distinct paralogs. We identify 57 additional regions covering 0.82% of the mouse genome with this property (**Figure 9** and **Supplementary Table 6**). These regions have gene density similar to unique sequence and are strongly enriched for genes involved in odorant sensing, pheromone recognition and immunity that play important roles in social behavior and speciation (Hurst *et al*. 2001). Excess polymorphism at these loci in mouse has previously been attributed to some combination of incomplete lineage sorting and diversifying selection (White *et al*. 2009; Keane *et al*. 2011). Our results suggest that inferences regarding the strength of selection on highly polymorphic loci in regions of subject to recurrent duplication and loss should be treated with caution.

Accurate deconvolution of recent duplications remains a difficult task that requires painstaking manual effort. Clone-based and/or single-molecule long-read sequencing remain the gold standard techniques. But short reads at sufficient depth nonetheless contain a great deal of information. We exploited the specific properties of *R2d2* in the WSB/EiJ mouse strain — many highly-similar copies of *R2d2* relative to the single divergent *R2d1* copy — to obtain a nearly complete assembly of *R2d2* from short reads (**Supplementary Figure 8**). With the sequence of both the *R2d1* and *R2d2* paralogs in hand, we were able to recognize several remarkable features of *R2d2* that are discussed in detail below.

### Long-tract gene conversion

Previous studies of non-allelic gene conversion in mouse and human have focused either on relatively small (<5 kbp) intervals within species, or have applied phylogenetic methods to multiple paralogs from a single reference genome (Dumont and Eichler 2013). This study is the first, to our knowledge, with the power to resolve large (>5 kbp) non-allelic gene conversion events on an autosome in a population sample. We identify conversion tracts up to 119 kbp in length, orders of magnitude longer than tracts arising from allelic conversion events during meiosis. Gene conversion at this scale can rapidly and dramatically alter paralogous sequences, including — as shown in **Figure 5** — the sequences of essential protein-coding genes. This process has been implicated as a source of disease alleles in humans (Chen *et al*. 2007).

Importantly, we were able to identify non-allelic exchanges in *R2d1* as such only because we were aware of the existence of *R2d2* in other lineages. In this case the transfer of paralogous *R2d2* sequence into *R2d1* creates the appearance of deep coalescence among *R2d1* sequences. Ignoring the effect of gene conversion would cause us to overestimate the degree of polymorphism at *R2d1* by an order of magnitude, and would bias any related estimates of population-genetic parameters (for instance, of effective population size).

Our data are not sufficient to estimate the rate of non-allelic gene conversion between *R2d2* and homologous loci. At minimum we have observed two distinct events: one from *R2d2* into *R2d1*, and a second from *R2d2* into retro-*Cwc22*. From a single conversion event replacing most of *R2d1* with *R2d2*-like sequence, the remaining shorter conversion tracts could be generated by recombination with *R2d1* sequences. Because we find converted haplotypes in both *M. m. musculus* and *M. m. domesticus*, the single conversion event would have had to occur prior to the divergence of the three *M. musculus* subspecies and subsequently remain polymorphic in the diverged populations. We note that all conversion tracts we observed are polarized: *R2d2* is always the donor.

The other possibility is that non-allelic gene conversion between *R2d* sequences is recurrent. Recurrent gene conversion homogenizes duplicate sequences, coupling their evolutionary trajectories (“concerted evolution”, Dover 1982). The absolute sequence divergence (~2%) between *R2d1* and *R2d2* (**Figure 1B**) argues against a hypothesis of ongoing gene conversion between these loci. However, we cannot rule out a role for gene conversion in maintaining sequence identity between multiple copies of *R2d* located in *R2d2*. This would help explain the reduced diversity within *R2d2* versus *R2d1* (**Figure 8C**). For sequences with copy number greater than two – such as the *R2d2* casssette in *R2d2*^*HC*^ alleles – gene conversion tends to slow the accumulation of new mutations in a copy-number dependent manner (Nagylaki and Petes 1982). New mutations arising in any single copy are prone to loss not only by drift but also by being “pasted-over” by gene conversion from the intact copies which outnumber the mutant (Melamed and Kupiec 1992).

*R2d2* is not unlike the male-specific region of the Y chromosome in mouse (Soh *et al*. 2014) and human (Rozen *et al*. 2003). The large palindromic repeats on chrY are homogenized by frequent non-allelic gene conversion (Hallast *et al*. 2013) such that they have retained >99% sequence identity to each other even after millions of years of evolution. Frequent non-allelic gene conversion has also been documented in arrays of U2 snRNA genes in human (Liao 1997), and in rRNA gene clusters (Eickbush and Eickbush 2007) and centromeric sequences (Schindelhauer 2002; Shi *et al*. 2010) in several species.

### Pervasive copy-number variation

Clusters of segmental duplications have long been known to be hotspots of copy-number variation in populations (Bailey and Eichler 2006; She *et al*. 2008) and *de novo* mutations in pedigrees (Egan *et al*. 2007). Recent large-scale sequencing efforts have revealed the existence of thousands of multiallelic CNVs segregating in human populations (Handsaker *et al*. 2015).

We have surveyed *R2d2* copy number in a large and diverse sample of laboratory and wild mice, and have shown that it varies from 0 to >80 in certain *M. m. domesticus* populations (**Figure 8A**). In a cohort of outbred mice expected to be hemizygous for an *R2d2*^*HC*^ allele from WSB/EiJ (33 diploid copies) we estimate that large deletions, >2 Mbp in size, occur at a rate of 3.2% (95% bootstrap CI 1.1% — 6.0%) per generation. This estimate of the mutation rate for CNVs at *R2d2* should be regarded as a lower bound. The power of our copy-number assay to discriminate between copy numbers above ~25 is low, so that the assay is much more sensitive to losses than to gains. Even our lower-bound mutation rate exceeds that of the most common recurrent deletions in human (~1 per 7000 live births) (Turner *et al*. 2007) and is an order of magnitude higher than the most active CNV hotspots described to date in the mouse (Egan *et al*. 2007).

However, the structural mutation rate appears to depend strongly on the diplotype configuration at *R2d2*. As **Figure 1D** shows, individuals heterozygous for an *R2d2*^*HC*^ haplotype and an *R2d2*-null haplotype are in fact hemizygous for several megabases of DNA in *R2d2*. This has important consequences. High mutation rates are observed only in the context of populations in which hemizygosity for *R2d2*^*HC*^ is common (**Figure 7**): highest in the DO, and to a lesser extent in wild *M. m. domesticus* populations harboring both *R2d2*^*HC*^ and *R2d2*-null alleles. Homozygosity for *R2d2*^*HC*^ is not associated with mutability: in 8 recombinant inbred lines from the Collaborative Cross which are homozygous for an *R2d2*^*HC*^ haplotype, we observed zero new mutations in at least 400 meioses, through both the male and female germline (8 lines × 2 meioses/generation × 25 or more generations of inbreeding). Sex also appears to have a role in determining the mutation rate at *R2d2:* in a pedigree in which all females were hemizygous for *R2d2*^*HC*^, zero new mutations were observed in 1256 meioses (data not shown).

Taken together, these observations hint at a common structural or epigenetic mechanism affecting the resolution of double-strand breaks in large tracts of unpaired (*i.e*. hemizygous) DNA during male meiosis. At least one other study in mouse has hinted that hemizygous SDs on the sex chromosomes are unstable in inter-subspecific hybrids (Scavetta and Tautz 2010). Both the obligate-hemizygous sex chromosomes and large unpaired segments on autosomes are epigenetically marked for transcriptional silencing during male meiotic prophase (Laan 2004; Baarends *et al*. 2005), and are physically sequestered into a structure called the sex body. Repair of double-strand breaks within the sex body is delayed relative to the autosomes (Mahadevaiah *et al*. 2001) and involves a different suite of proteins (Turner *et al*. 2004). We hypothesize that these male-specific pathway(s) are generally error-prone in the presence of non-allelic homologous sequences.

### Origin and distribution of an allele subject to meiotic drive

Females heterozygous for a high-and low-copy allele at *R2d2* preferentially transmit the high-copy allele to progeny via meiotic drive (Didion *et al*. 2015). Meiotic drive can rapidly alter allele frequencies in laboratory and natural populations (Lindholm *et al*. 2016), and we recently showed that high-copy alleles of *R2d2* (*R2d2*^*HC*^) sweep through laboratory and natural populations despite reducing the fitness of heterozygous females (Didion *et al*. 2016). These “selfish sweeps” account for the marked reduction in within-population diversity in the vicinity of *R2d2* (Figure 8B).

The present study sheds additional light on the age, origins and fate of *R2d2*^*HC*^ alleles. We find that *R2d2*^*HC*^ alleles have a single origin in *M. m. domesticus*. They are present in several different “chromosomal races” — populations fixed for specific Robertsonian translocations between which gene flow is limited (Hauffe and Searle 1993) — indicating that they were likely present at intermediate frequency prior to the origin of the chromosomal races within the past 6,000 to 10,000 years (Nachman *et al*. 1994) and were dispersed through Europe as mice colonized the continent from the south and east (Boursot *et al*. 1993). The presence of *R2d2*^*HC*^ in a non-M. *m. domesticus* sample (SPRET/EiJ, *M. spretus* from Cadiz, Spain) is best explained by recent introgression following secondary contact with *M. m. domesticus* (Bonhomme *et al*. 2007; Yang *et al*. 2011).

### A new member of the *Cwc22* family

The duplication that gave rise to *R2d2* also created a new copy of *Cwc22*. Based on our assembly of the *R2d2* sequence, the open reading frame of *Cwc22*^*R2d2*^ is intact and encodes a nearly full-length predicted protein that retains the two key functional domains characteristic of the *Cwc22* family. Inspection of RNA-seq data from samples with high copy number at *R2d2* reveals several novel transcript isoforms whose expression appears to be copy-number‐ and tissue-dependent. In testis, the most abundant isoform retains an intron containing an ERV insertion (red arrow in **Figure 4**), consistent with the well-known transcriptional promiscuity in this tissue. The most abundant isoforms in adult brain is unusual in that its stop codon is in an internal exon which is followed by a 7 kbp 3’ UTR in the terminal exon. Transcripts with a stop codon in an internal exon are generally subject to nonsense-mediated decay (NMD) triggered by the presence of exon-junction complexes downstream the stop codon. Curiously, *Cwc22* is itself a member of the exon-junction complex (Steckelberg *et al*. 2012).

That an essential gene involved in such a central biochemical pathway should both escape NMD and be overexpressed more than tenfold is surprising. Preliminary data from the Diversity Outbred population shows that the *R2d2*^*HC*^ allele is associated with elevated levels of both *Cwc22* transcripts and protein in adult liver (Gary Churchill, personal communication). Further studies will be required to determine the distribution of transcription and translation of *Cwc22* across isoforms, tissues and developmental stages.

### Conclusions and future directions

Our detailed analysis of the evolutionary trajectory of *R2d2* provides insight into the fate of duplicated sequences over short (within-species) timescales. The exceptionally high mutation rate and low recombination associated specifically with hemizygous *R2d2*^*HC*^ alleles motivate hypotheses regarding the biochemical mechanisms which contribute to observed patterns of polymorphism at this and similar loci. Finally, the birth of a new member of the deeply conserved *Cwc22* gene family in *R2d2* provides an opportunity to test predictions regarding the evolution of young duplicate gene pairs.

## METHODS

### Mice

Wild *M. musculus* mice used in this study were trapped at a large number of sites across Europe, the United States, the Middle East, northern India and Taiwan (Figure 8A). Trapping was carried out in accordance with local regulations and with the approval of all relevant regulatory bodies for each locality and institution. Trapping locations are listed in **Supplementary Table 1**. Most samples have been previously published (Didion *et al*. 2016).

Tissue samples from the progenitors of the wild-derived inbred strains ZALENDE/EiJ (*M. m. domesticus*), TIRANO/EiJ (*M. m. domesticus*) and SPRET/EiJ (*M. spretus*) were provided by Muriel Davisson, as described in Didion *et al*. (2016).

Tissue samples from the high running (HR) selection and intercross lines were obtained as described in Didion *et al*. (2016).

Female Diversity Outbred mice used for estimating mutation rates at *R2d2* were obtained from the Jackson Laboratory and housed with a single FVB/NJ male. Progeny were sacrificed at birth by cervical dislocation in order to obtain tissue for genotyping.

All live laboratory mice were handled in accordance with the IACUC protocols of the University of North Carolina at Chapel Hill.

### DNA preparation

*High molecular weight DNA*. High molecular weight DNA was obtained for samples genotyped with the Mouse Diversity Array or subject to whole-genome sequencing. Genomic DNA was extracted from tail, liver or spleen using a standard phenol-chloroform procedure (Sambrook and Russell 2006). High molecular weight DNA for most inbred strains was obtained from the Jackson Laboratory, and the remainder as a generous gift from Francois Bonhomme and the University of Montpellier Wild Mouse Genetic Repository.

*Low molecular weight DNA*. Low molecular weight DNA was obtained for samples to be genotyped on the MegaMUGA array (see “Microarray genotyping” below). Genomic DNA was isolated from tail, liver, muscle or spleen using Qiagen Gentra Puregene or DNeasy Blood & Tissue kits according to the manufacturer's instructions.

### Whole-genome sequencing and variant discovery

*Inbred strains*. Sequencing data for inbred strains of mice except ZALENDE/EiJ and LEWES/EiJ was obtained from the Sanger Mouse Genomes Project website (ftp://ftp-mouse.sanger.ac.uk/current_bams) as aligned BAM files. Details of the sequencing pipeline are given in Keane *et al*. (2011). Coverage ranged from approximately 25X to 50X per sample.

The strains LEWES/EiJ and ZALENDE/EiJ were sequenced at the University of North Carolina High-Throughput Sequencing Facility. Libraries were prepared from high molecular weight DNA using the Illumina TruSeq kit and insert size approximately 250 bp, and 2x100bp paired-end reads were generated on an Illumina HiSeq 2000 instrument. LEWES/EiJ was sequenced to approximately 12X coverage and ZALENDE/EiJ to approximately 20X. Alignment was performed as in Keane *et al*. (2011).

*Wild mice*. Whole-genome sequencing data from 26 wild *M. m. domesticus* individuals described in Pezer *et al*. (2015) was downloaded from ENA under accession #PRJEB9450. Coverage ranged from approximately 12X to 20X per sample. An additional two wild *M. m. domesticus* individuals, IT175 and ES446, were sequenced at the University of North Carolina to approximate coverage 8X each. Raw reads from an additional 10 wild *M. m. castaneus* described in Halligan *et al*. (2013), sequenced to approximately 20X each, were downloaded from ENA under accession #PRJEB2176. Reads for a single *Mus caroli* individual sequenced to approximately 40X were obtained from ENA under accession #PRJEB2188. Reads for each sample were realigned to the mm10 reference using bwa-mem v0.7.12 with default parameters (Li 2013). Optical duplicates were removed with samblaster (Faust and Hall 2014). *Variant discovery*. Polymorphic sites on chromosome 2 in the vicinity of *R2d2* (**Figure 8B**) were called using freebayes v0.9.21-19-gc003c1e (Garrison and Marth 2012) with parameters “— standard-filters” using the Sanger Mouse Genomes Project VCF files as a list of known sites (parameter “ — @”). Raw calls were filtered to have quality score > 30, root mean square mapping quality > 20 (for both reference and alternate allele calls) and at most 2 alternate alleles.

### Copy-number estimation

*R2d* copy number was estimated using qPCR as described in Didion *et al*. (2016). Briefly, we used commercial TaqMan assays against intron-exon boundaries in *Cwc22* (Life Technologies assay numbers Mm00644079_cn and Mm00053048_cn) to determine copy number relative to reference genes *Tert* (cat. no. 4458368, for target Mm00644079_cn) or *Tfrc* (cat. no. 4458366, for target Mm00053048_cn). Cycle thresholds for *Cwc22* relative to the reference gene were normalized across assay batches using linear mixed models with batch and target-reference pair treated as random effects. Control samples with known haploid *R2d* copy numbers of 1 (C57BL/6J), 2 (CAST/EiJ), 17 (WSB/EiJxC57BL/6J)F1 and 34 (WSB/EiJ) were included in each batch.

Samples were classified as having 1, 2 or >2 haploid copies of *R2d* using linear discriminant analysis. The classifier was trained on the normalized cycle thresholds of the control samples from each plate, whose precise integer copy number is known, and applied to the remaining samples.

### Microarray genotyping

Genome-wide genotyping was performed using MegaMUGA, the second version of the Mouse Universal Genotyping Array platform (Neogen/GeneSeek, Lincoln, NE) (Morgan *et al*. 2016). Genotypes were called using the GenCall algorithm implemented in the Illumina BeadStudio software (Illumina Inc, Carlsbad, CA). For quality control we computed, for each marker *i* on the array: $S_i = *X*_*i*_ + *Y*_*i*_, where *X*_*i*_ and *Y*_*i*_ are the normalized hybR2d2zation intensities for the two alleles. The expected distribution of Si was computed from a large set of reference samples. We excluded arrays for which the distribution of *S*_*i*_ was substantially shifted from this reference; in practice, failed arrays can be trivially identified in this manner (Morgan *et al*. 2016). Access to MegaMUGA genotypes was provided by partnership between the McMillan and Pardo-Manuel de Villena labs and the UNC Systems Genetics Core Facility.

Additional genotypes for inbred strains and wild mice from the Mouse Diversity Array were obtained from Yang *et al*. (2011).

### *De novo* assembly of *R2d2*

Raw whole-genome sequencing reads for WSB/EiJ from the Sanger Mouse Genomes Project were converted to a multi-string Burrows-Wheeler transform and associated FM-index (msBWT) (Holt and McMillan 2014) using the msbwt v0.1.4 Python package (https://pypi.python.org/pypi/msbwt). The msBWT and FM-index implicitly represent a suffix array of sequencing to provide efficient queries over arbitrarily large string sets. Given a seed *k*-mer present in that string set, this property can be exploited to rapidly construct a de Bruijn graph which can in turn be used for local *de novo* assembly of a target sequence (**Supplementary Figure 8A**). The edges in that graph can be assigned a weight (corresponding to the number of reads containing the *k* + 1-mer implied by the edge) which can be used to evaluate candidate paths when the graph branches (**Supplementary Figure 8B**).

*R2d2* was seeded with the 30 bp sequence (TCTAGAGCATGAGCCTCATTTATCATGCCT) at the proximal boundary of *R2d1* in the GRCm38/mm10 reference genome. A single linear contig was assembled by “walking” through the local de Bruijn graph. Because WSB/EiJ has ~33 copies of *R2d2* and a single copy of *R2d1*, any branch point in the graph which represents a paralogous variant should having outgoing edges with weights differing by a factor of approximately 33. Furthermore, when two (or more) branch points occur within less than the length of a read, it should be possible to “phase” the underlying variants by following single reads through both branch points (**Supplementary Figure 8B**). We used these heuristics to assemble the sequence of *R2d2* (corresponding to the higher-weight path through the graph) specifically.

After assembling a chunk of approximately 500 bp the contig was checked for colinearity with the reference sequence (*R2d1*) using BLAT and CLUSTAL-W2 (using the EMBL-EBI web server: http://www.ebi.ac.uk/Tools/msa/clustalw2/).

Repetitive elements such as retroviruses are refractory to assembly with our method. Upon traversing into a repetitive element, the total edge weight (total number of reads) and number of branch points (representing possible linear assembled sequences) in the graph become large. It was sometimes possible to assemble a fragment of a repetitive element at its junction with unique sequence but not to assemble unambiguously across the repeat. Regions of unassembleable sequence were marked with blocks of Ns, and assembly re-seeded using a nearby k-mer from the reference sequence. The final contig is provided in FASTA format in **Supplementary File 1**.

The final contig was checked against its source msBWT by confirming that each 30-mer in the contig which did not contain an N was supported by at least 60 reads. A total of 16 additional haplotypes in 8 regions of *R2d* totaling 16.9 kbp (**Supplementary Table 6**) were assembled in a similar fashion, using the WSB *R2d2* contig and the *R2d1* reference sequence as guides. Multiple sequence alignments from these regions are provided in **Supplementary File 1**.

### Sequence analysis of *R2d2* contig

*Pairwise alignment of* R2d *paralogs*. The reference *R2d2* sequence and our *R2d1* contig were aligned using LASTZ v1.03.54 (http://www.bx.psu.edu/~rsharris/lastz/) with parameters “ –step=10 –seed=match12 – notransition –exact=20 –notrim –identity=95”.

*Transposable element (TE) content*. The *R2d2* contig was screened for TE insertions using the RepeatMasker web server (http://www.repeatmasker.org/cgi-bin/WEBRepeatMasker) with species set to “mouse” and default settings otherwise. As noted previously, we could not assemble full-length repeats, but the fragments we could assemble at junctions with unique sequence allowed identification of some candidate TEs to the family level. *R2d1*-specific TEs were defined as TEs annotated in the RepeatMasker track at the UCSC Genome Browser with no evidence (no homologous sequence, and no Ns) at the corresponding position in the *R2d2* contig. Candidate *R2d2*-specific TEs were defined as gaps >= 100 bp in size in the alignment to *R2d1* for which the corresponding *R2d2* sequence was flagged by RepeatMasker.

*Gene conversion tracts*. To unambiguously define gene conversion events without confounding from paralogous sequence, we examined 15 wild *M. m. domesticus* samples and 37 laboratory strains with evidence of 2 diploid copies of *R2d*. We confirmed that these copies of *R2d* were located at *R2d1* by finding read pairs spanning the junction between *R2d1* and neighboring sequence. Gene conversion tracts were delineated as clusters of derived alleles shared with *R2d2*. Using a pairwise of alignment of *R2d2* and *R2d1* we identified single-nucleotide variants between the two sequences, and queried those sites in aligned reads for *Mus caroli*. If the *Mus caroli* and *R2d1* shared an allele, we recorded the site as a derived allele informative for the presence of *R2d2*. We used the resulting list of 1,411 informative sites to query aligned reads for the samples of interest and recorded, for each site and each sample, whether the derived allele (*R2d2*), ancestral allele (*R2d1*) or both alleles were present. Conversion tracts were then identified by manual inspection. Boundaries of conversion tracts were defined at approximately the midpoint between the first *R2d1*- (or *R2d2*-) specific variant and the last *R2d2*- (or *R2d1*-) specific variant.

*Sequence diversity in* R2d1 *and* R2d2. Assembling individual copies of *R2d2* is infeasible in high-copy samples. Instead we treated each *R2d* unit as an independent sequence and used the number of segregating sites to estimate sequence diversity. Segregating sites were defined as positions in a collection of alignments (BAM files) with evidence of an alternate allele. To identify segregating sites we used freebayes v0.9.21-19-gc003c1e (Garrison and Marth 2012) with parameters “-ui-Kp 20 —use-best-n-alleles 2-m 8”. These parameters treat each sample as having ploidy up to 20, impose an uninformative prior on genotype frequencies, and limit the algorithm to the discovery of atomic variants (SNVs or short indels, not multinucleotide polymorphisms or other complex events) with at most 2 alleles at each segregating site. Sites in low-complexity sequence (defined as Shannon entropy < 1.6 in the 30 bp window centered on the site) or within 10 bp of another variant site were further masked, to minimize spurious calls due to ambiguous alignment of indels. To avoid confounding with the retrocopies of *Cwc22* outside *R2d*, coding exons of *Cwc22* were masked. Finally, sites corresponding to an unaligned or gap position in the pairwise alignment between *R2d1* and *R2d2* were masked.

To compute diversity in *R2d1* we counted segregating sites in 12 wild *M. m. domesticus* samples with 2 diploid copies of *R2d* (total of 24 sequences), confirmed to be in *R2d2* by the presence of read pairs spanning the junction between *R2d1* and neighboring sequence. To compute diversity in *R2d2*, we counted segregating sites in 14 wild *M. m. domesticus* samples with >2 diploid copies of *R2d* (range 3 — 83 per sample; total of 406 sequences) but excluded sites corresponding to variants among *R2d1* sequences. Remaining sites were phased to *R2d2* by checking for the presence of a 31-mer containing the site and the nearest *R2d1-vs-R2d2* difference in the raw reads for each sample using the corresponding msBWT. Sequence diversity was then computed using Watterson’s estimator (Watterson 1975), dividing by the number of alignable bases (128973) to yield a per-site estimate. Standard errors were estimated by 100 rounds of resampling over the columns in the *R2d1*-vs-*R2d2* alignment.

### Analyses of *Cwc22* expression

*RNA-seq read alignment*. Expression of *Cwc22* was examined in adult whole brain using data from Crowley *et al*. (2015), SRA accession #SRP056236. Paired-end reads (2x100bp) were obtained from 8 replicates each of 3 inbred strains: CAST/EiJ, PWK/PhJ and WSB/EiJ. Raw reads were aligned to the mm10 reference using STAR v2.4.2a (Dobin *et al*. 2012) with default parameters for paired-end reads. Alignments were merged into a single file per strain for further analysis. Expression in adult testis was examined in 23 wild-derived inbred strains from Phifer-Rixey *et al*. (2014) SRA accession #PRJNA252743. Single-end reads (76bp) were aligned to the mm10 genome with STAR using default parameters for single-end, non-strand-specific reads.

*Transcript assembly*. Read alignments were manually inspected to assess support for *Cwc22* isoforms in Ensembl v83 annotation. To identify novel isoforms in *R2d2*, we applied the Trinity v0.2.6 pipeline (Grabherr *et al*. 2011) to the subset of reads from WSB/EiJ which could be aligned to *R2d1* plus their mates (a set which represents a mixture of *Cwc22*^*R2d1*^ and *Cwc22*^*R2d2*^ reads). De novo transcripts were aligned both to the mm10 reference and to the *R2d2* contig using BLAT, and were assigned to *R2d1* or *R2d2* based on sequence similarity. Because expression from *R2d2* is high in WSB/EiJ, *R2d2*-derived transcripts dominated the assembled set. Both manual inspection and the Trinity assembly indicated the presence of retained introns and an extra 3’ exon, as described in the Results. To obtain a full set of Cwc22 transcripts including those of both *R2d1* and *R2d2* origin, we supplemented the *Cwc22* transcripts in Ensembl v83 with their paralogs from *R2d2* as determined by a strict BLAT search against the *R2d2* contig. We manually created additional transcripts reflecting intron-retention and 3’ extension events described above, and obtained their sequence from the *R2d2* contig.

*Abundance estimation*. Relative abundance of *Cwc22* paralogs was estimated using kallisto v0.42.3 (Bray *et al*. 2015) with parameters “—bias” (to estimate and correct library-specific sequence-composition biases). The transcript index used for pseudoalignment and quantification included only the *Cwc22* targets.

### Phylogenetic analyses

*Tree for* R2d. Multiple sequence alignments for 8 the regions in **Supplementary Figure 5** were generated using MUSCLE (Edgar 2004) with default parameters. The resulting alignments were manually trimmed and consecutive gaps removed. Phylogenetic trees were inferred with RAxML v8.1.9 (Stamatakis 2014) using the GTR+gamma model with 4 rate categories and *M. caroli* as an outgroup. Uncertainty of tree topologies was evaluated using 100 bootstrap replicates.

*Divergence time*. The time of the split between *R2d1* and *R2d2* was estimated using the Bayesian method implemented in BEAST v1.8.1r6542 (Drummond *et al*. 2012). We assumed a divergence time for *M. caroli* of 5 Mya and a strict molecular clock, and analyzed the concatenated alignment for our *de novo* assembled regions under the GTR+gamma model with 4 rate categories and allowance for a proportion of invariant sites. The chain was run for 10 million iterations with trees sampled every 1000 iterations.

*Local phylogeny around* R2d2. Genotypes for 173 SNPs in the region surrounding *R2d2* (chr2: 83 — 84 Mb) were obtained for 90 individuals representing both laboratory and wild mice genotyped with the Mouse Diversity Array (Yang *et al*. 2011) (**Supplementary Table 1**). Individuals with evidence of heterozygosity (>3 heterozygous calls) were excluded to avoid ambiguity in phylogenetic inference. A distance matrix for the remaining 62 samples was created by computing the proportion of alleles shared identical by state between each pair of samples. A neighbor-joining tree (**Figure 3C**) was inferred from the distance matrix and rooted at the most recent common ancestor of the *M. musculus*- and non-M. *musculus* samples.

Cwc22 *coding sequences*. To create the tree of Cwc22 coding sequences, we first obtained the sequences of all its paralogs in mouse. The coding sequence of *Cwc22*^*R2d1*^ (RefSeq transcript NM_030560.5) was obtained from the UCSC Genome Browser and aligned to our *R2d2* contig with BLAT to extract the exons of *Cwc22*^*R2d2*^. The coding sequence of retro-*Cwc22* (genomic sequence corresponding to GenBank cDNA AK145290) was obtained from the UCSC Genome Browser. Coding and protein sequences of *Cwc22* homologs from non-M. *musculus* species were obtained from Ensembl (Cunningham *et al*. 2014). The sequences were aligned with MUSCLE and manually trimmed, and a phylogenetic tree estimated as described above.

We observed that the branches in the rodent clade of the *Cwc22* tree appeared to be longer than branches for other taxa. We used PAML (Yang *et al*. 2007) to test the hypothesis that *Cwc22* is under relaxed purifying selection in rodents using the branch-site model (null model “model = 2, NSsites = 2, fix_omega = 1”; alternative model “model = 2, NSsites = 2, omega = 1, fix_omega = 1”) as described in the PAML manual. This is a test of difference in evolutionary rate on a “foreground” branch (*ω*_1_) — in our case, the rodent clade — relative to the tree-wide “background” rate (*ω*_0_). The distribution of the test statistic is an even mixture of a *χ*^2^ distribution with 1 df and a point mass at zero; to obtain the p-value, we calculated the quantile of the *χ*^2^ distribution with 1 df and divided by 2.

### Genome-wide sequence divergence

The msBWT of a collection of whole-genome sequencing reads can be used to estimate the divergence between the corresponding template sequence (*i.e*. genome) and a reference sequence as follows. Non-overlapping *k*-mers from the reference sequence are queried against the msBWT. (The value of *k* is chosen such that nearly all *k*-mers drawn from genomic sequence exclusive of repetitive elements.) Let *x* be the count of reads containing an exact match to the *k*-mer or its reverse complement. If the template and reference sequence are identical, standard theory for shotgun sequencing (Lander and Waterman 1988) holds that *P*(*x* > 0|*λ*) = 1 – *e*^−λ^, where *λ* is the average sequencing coverage. We assume *P*(*x* > 0|*λ*) ≈ 1, which is satisfied in practice for high-coverage sequencing.

However, if a haploid template sequence contains at least one variant (versus the reference) within the queried *k*-mer, it will be the case that *x* = 0. We use this fact and assume that mutations arise along a sequence via a Poisson process to estimate the rate parameter *α* from the proportion of *k*-mers that have read count zero. Let *m* be the number of mutations arising between a target and reference in a window of length *L*, and *y* the number of *k*-mers in that window with nonzero read count. Then *P*(*m* = 0|*α*) = *e*^*-αL*^ and a simple estimator for 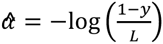.

Interpretation of *α* is straightforward in the haploid case: it is the per-base rate of sequence divergence between the template sequence and the reference sequence. In the diploid case it represents a lower bound on the sequence divergence of the two homologous chromosomes.

We applied this estimator with *k* = 31 and *L* = 1000×*k* = 31 kbp to msBWTs for 7 inbred strains (3 *M. m. domesticus*, 1 *M. m. musculus*, 1 *M. m. castaneus*, 1 *M. spretus*, 1 *M. caroli*) and 2 wild *M. m. domesticus* individuals (IT175, ES446) using the GRCm38/mm10 mouse reference sequence as the source of *k*-mer queries. As shown in **Figure 9A**, the mode of the distribution of divergence values matches what is expected based on the ancestry of the samples with respect to the reference. To identify divergent regions, we fit a discrete-time hidden Markov model (HMM) to the windows divergence values. The HMM had two hidden states: “normal” sequence, with emission distribution *N*(0.005,0.005) and initial probability 0.99; and “divergent” sequence, with emission distribution *N*(0.02,0.005) and initial probability 0.01. The transmission probability between states was 1×10^−5^. Posterior decodings were obtained via the Viterbi algorithm, as implemented in the R package HiddenMarkov (https://cran.r-project.org/package=HiddenMarkov).

Significance tests for overlap with genomic features were performed using the resampling algorithm implemented in the Genomic Association Tester (GAT) package for Python (https://pypi.python.org/pypi/gat). Segmental duplications were obtained from the genomicSuperDups table of the UCSC Genome Browser and genes from Ensembl v83 annotation.

### Analyses of recombination rate at *R2d2*

To test the effect of *R2d2* copy number on local recombination rate examined recombination events accumulated during the first 16 generations of breeding of the Diversity Outbred (DO) population, in which the high-copy *R2d2* allele from WSB/EiJ is segregating. Founder haplotype reconstructions were obtained for 4,640 DO individuals reported in (Didion *et al*. 2016), and recombination events were identified as junctions between founder haplotypes. We compared the frequency of junctions involving a WSB/EiJ haplotype to junctions not involving a WSB/EiJ haplotype over the region chr2: 75-90 Mb. Within each generation we also tested for differences in the lengths of haplotype blocks overlapping *R2d2* using one-sided Wilcoxon rank-sum tests (alternative hypothesis: WSB/EiJ haplotypes longer than others). Resulting *p*-values were subject to Bonferroni correction: for nominal significance level *α* = 0.01, the corrected threshold is 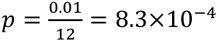.

We also estimated the difference between observed and expected recombination fraction in 11 experimental crosses in which one of the parental lines was segregating for a high-copy allele at *R2d2*. We obtained expected recombination fractions from the standard mouse genetic map (Cox *et al*. 2009), which was constructed from crosses between strains lacking *R2d2*^*HC*^ alleles. Genotype data was obtained from The Jackson Laboratory’s Mouse Phenome Database QTL Archive (http://phenome.jax.org/db/q?rtn=qtl/home). Recombination fractions were calculated using R/qtl (http://rqtl.org/). Confidence intervals for difference between observed and expected recombination fractions were calculated by 100 iterations of nonparametric bootstrapping over individuals in each dataset.

Results of these analyses are presented in **Supplementary Figure 7**.

## Data availability

All *de novo* assemblies used in this study are included in **Supplementary File 1**. The data structures on which the assemblies are based, and the interactive computational tools used for assembly, are publicly available at http://www.csbio.unc.edu/CEGSseq/index.py?run=MsbwtTools. Genotype data used for mapping the location of R2d2 and defining associated haplotypes (Figure 3B–C) are available on Dryad (accession XXX).

## Acknowledgments

We thank all the scientists and personnel who collected and processed the wild mouse samples used in this study. In particular we thank Francois Bonhomme for providing samples from wild-derived inbred strains housed at the University of Montpellier Wild Mouse Genetic Repository, and Ted Garland for providing tissue samples from the HR selection lines and related crosses. This work was supported by National Institutes of Health grants P50GM076468 (FPMdV), U19AI100625 (FPMdV, APM), F30MH103925 (APM), T32GM067553 (JPD, APM), and by Vaadia-BARD Postdoctoral Fellowship Award FI-12 478-13 to LY. Additional support was provided by Cancer Research UK, the European Research Council, EMBO Young Investigator Programme (DTO), European Molecular Biology Laboratory (DTO, PF) and the Wellcome Trust (WT095908) (PF) and (WT098051) (PF, DTO); and finally by the European Union's Seventh Framework Programme (FP7/2007-2013) under grant agreement HEALTH-F4-2010-241504 (EURATRANS).

## Supplementary material

**Supplementary Figure 1**. Conservation of synteny between mouse and four other mammals around *Cwc22*^*R2d1*^ (upper panel) indicates that the *R2d1* sequence remains in its ancestral position. Chevrons represent genes, alternating white and grey, and are oriented according to the strand on which the gene is encoded. *Cwc22*^*R2d2*^ is novel in the mouse but its position relative to genes with conserved order is shown in the lower panel. Note that synteny is disrupted in mouse and rat distal to *R2d2*.

**Supplementary Figure 2.** Pairwise alignment of *R2d2* contig (top) to the *R2d1* reference sequence (bottom). Dark boxes show position of repetitive elements present in both sequences; syntenic positions are connected by grey anchors, and blank space represents aligned bases in both sequences. Orange boxes indicate position of repetitive elements present in the *R2d1* sequence but not detected in *R2d2;* blue boxes indicate position of elements in *R2d2* but not *R2d1. Cwc22* transcripts are shown below the alignment.

**Supplementary Figure 3.** Phylogenetic tree constructed from amino acid sequences for mammalian *Cwc22* homologs (including all three mouse paralogs) with chicken as an outgroup. Node labels indicate support in 100 bootstrap replicates.

**Supplementary Figure 4**. Alignment of amino acid sequences from mouse *Cwc22*^*R1d1*^, *Cwc22*^*R2d2*^ and retro-*Cwc22*, plus *Cwc22* orthologs from 19 other placental mammals plus opossum, platypus and chicken as outgroups. Residues are colored according to biochemical properties and gaps are shown in grey. Information content of each column in the alignment, measured as the Jenson-Shannon divergence, is plotted in the lower panel.

**Supplementary Figure 5**. Diagnostic variants used to identify gene conversion tracts between *R2d2* and *R2d1*. Each column represents a single variant (total of 1,411) between *R2d1* and *R2d2* for which *R2d2* has the derived allele, and each row a single individual with whole-genome sequence data available (named in **Supplementary Table 1**). Points are colored according to the “allelotype” of each variant detected in 8 or more reads in each sample: 0:0 (red), neither *R2d1* nor *R2d2* allele present; 1:0, *R2d1* but not *R2d2* (orange); 1:1, both *R2d1* and *R2d2* (green); 0:1, *R2d2* but not *R2d1* (blue). Individuals are grouped according to diploid *R2d* copy number.

**Supplementary Figure 6**. Partial loss of *R2d2* with structural rearrangement. **(A)** Inferred structure of the *R2d1-R2d2* region in IR:AHZ_STND:015, a wild *M. m. domesticus* individual from Iran. *R2d1* is present on both chromosomes but only a fragment of *R2d2* remains on one chromosome, and it has been transposed into the retro-Cwc22 array. **(B)** Normalized depth of coverage (2 = normal diploid level) across *R2d*. Regions in grey represent reads from *R2d1* alone, while region in black captures reads from *R2d1* and *R2d2*, as shown by arrows from panel A. **(C)** Position of read pairs (red; not drawn to scale) with soft-clipped alignments to *R2d1*. The proximal read aligns in the 3’ UTR of *Cwc22*, and the distal read across an exon-intron boundary within the gene body. Note the “outward”-facing direction of the alignments. **(D)** Positions of the mates of the reads in panel C. Note that the x-axis is reversed so that the exons of retro-*Cwc22* (encoded on the plus strand) parallel those of *Cwc22* (encoded on the minus strand). The 3’ read maps across the boundary of th 3’ UTR of *Cwc22* and the ERV mediating the retrotransposition event. The 5’ read maps across two exon-exon boundaries in retro-*Cwc22*, so there is no ambiguity regarding its alignment to the retro-transposed copy. (E) Inferred structure of *Cwc22* paralogs in this sample. Note that one of the copies of retro-*Cwc22* is now a mosaic of retrotransposed and *Cwc22*^*R2d2_*^ derived sequence.

**Supplementary Figure 7**. Difference between expected and observed recombination fraction between markers flanking *R2d2* in experimental crosses in which at least one parent is segregating for a high-copy allele of *R2d2*. Thick and thin vertical bars show 90% and 95% confidence bounds, respectively, obtained by non-parametric bootstrap.

**Supplementary Figure 8. Targeted *de novo* assembly using the multi-string Burrows-Wheeler Transform (msBWT). (A)** The msBWT and its associated FM-index implicitly represent a suffix array of sequencing reads, such that read suffixes sharing a *k*-mer prefix are adjacent in the data structure. This allows rapid construction of a local de Bruijn graph starting from a *k*-mer seed (dark blue) and extending by successive *k*-mers (light blue) containing the (*k* — 1)-length suffix of the previous *k*-mer. A (*k* — 1)-length prefix with more than one possible suffix (red and orange) creates a branch point. Adjacent nodes in the graph with in-degree and out-degree one can be collapsed into a single node, yielding a simplified graph, which can then be traversed to obtain linear contig(s). **(B)** Paralogs of *R2d* can be disentangled using the local de Bruijn graph by exploiting differences in copy number. Edges in the graph are weighted by read count, and linear contigs for the *R2d1* and *R2d2* paralogs obtained by traversing the graph in a manner that minimizes the variance in edge weights along possible paths. Phase-informative reads (those overlapping multiple paralogous variants) provide a second source of evidence.

**Supplementary Table 1**. List of mouse samples used in this study, with their taxonomic designation, geographic origin, karyotype (STND, standard: 2n = 40; all others chromosomal races with Robertsonian translocations) and *R2d2* copy-number classification.

**Supplementary Table 2**. Transposable-element insertions private to *R2d1* or *R2d2*. Coordinates are offsets with respect to the start position of *R2d* (for *R2d1:* chr2: 77,869,657 in the reference genome; for *R2d2:* the beginning of the *de novo* assembled contig in **Supplementary File 1**.)

**Supplementary Table 3**. Frequency table of copy-number status by geographic origin for wild-caught and wild-derived Mus musculus individuals used in this study, stratified by subspecies. “Europe/Mediterranean” includes continental Europe, the United Kingdom and countries in the Mediterranean basin (Tunisia, Cyprus, Israel). “Asia” includes Asia, the Middle East and countries in the Indian Ocean basin (Madagascar).

**Supplementary Table 4**. Individuals from the Diversity Outbred population carrying *de novo* copy-number mutations at *R2d2*. Each was expected to be heterozygous for the WSB/EiJ allele (33 haploid copies).

**Supplementary Table 5**. Regions of excess divergence between wild or wild-derived mice and the mouse reference genome (GRCm38/mm10 build).

**Supplementary Table 6**. Regions of *R2d* targeted for *de novo* assembly in inbred strains.

**Supplementary File 1**. Compressed archive containining *R2d2* contig (from WSB/EiJ) and multiple sequence alignments from selected regions in **Supplementary Table 6**.

